# Continuous sensing of nutrients and growth factors by the mTORC1-TFEB axis

**DOI:** 10.1101/2021.08.07.455512

**Authors:** Breanne Sparta, Michael Pargett, Nont Kosaisawe, John G. Albeck

## Abstract

mTORC1 senses nutrient and growth factor status and phosphorylates downstream targets, including the transcription factor TFEB, to coordinate metabolic supply and demand. The molecular mechanisms of mTORC1 activation are thought to enforce a strict requirement for simultaneous amino acid and growth factor stimuli, but this model has not been evaluated with quantitative or single-cell methods. Here, we develop a series of fluorescent protein-TFEB fusions and investigate how combinations of stimuli jointly regulate signaling from mTORC1 to TFEB at the single-cell level. Live-cell imaging of individual cells revealed that mTORC1-TFEB signaling responds with graded changes to individual amino acid and growth factor inputs, rather than behaving as a logical “AND” gate. We find that mTORC1 inputs can be sequentially sensed, with responses that vary between mTORC1 substrates and are amplified by input from other kinases, including GSK3β. In physiologically relevant concentrations of amino acids, we observe fluctuations in mTORC1-TFEB signaling that indicate continuous responsiveness to nutrient availability. Our results clarify how the molecular regulation of mTORC1 enables homeostatic processes at the cellular level and provide a more precise understanding of its behavior as an integrator of multiple inputs.

## Introduction

To adapt to continuously changing microenvironments, cells must perceive the availability of multiple nutrients and adjust anabolic and catabolic processes accordingly. The kinase mTOR (mammalian Target Of Rapamycin) is the core of a regulatory network for sensing growth factors and metabolites, including amino acids, glucose, and cellular energy levels (ATP/ADP/AMP ratios). Collectively, these factors regulate the assembly and activity of mTOR Complex 1 (mTORC1), a heteromer of mTOR, Raptor, mLST8, PRAS40, and DEPTOR (Condon and Sabatini, 2019). Through both transcriptional control and direct protein modification, the kinase activity of this complex coordinates many downstream processes, including protein, nucleotide, and lipid synthesis (Shimobayashi and Hall, 2014). mTORC1 also regulates autophagy, the process of nutrient salvaging and organelle biogenesis, via the kinase ULK1 and the transcription factor TFEB (Kim et al., 2011; Settembre et al., 2011).

While mTORC1 regulates many essential processes in the cell, its activity is not required to sustain these processes at basal rates (Valvezan and Manning, 2019). Cell lines lacking mTORC1 function can be derived and propagated under *in vitro* conditions, where nutrients are in excess (Cybulski et al., 2012). However, mice die very early in embryogenesis if they are either deficient for mTORC1 (Guertin et al., 2006) or unable to inactivate mTORC1 under low amino acid conditions (Efeyan et al., 2013), indicating that mTORC1 regulation is needed under physiological conditions, where nutrient availability varies. These observations suggest that the essential physiological role of mTORC1 is to act as a “controller” that aligns the rates of catabolic and anabolic processes, with similarities to an engineered homeostatic control loop. However, our understanding of this controller function is unsatisfying from a quantitative perspective. Extensive work in homeostatic control across many fields has shown that understanding a controller’s function requires extensive characterization of its quantitative response across its full range of operating conditions. While many molecular details of mTORC1 activation have been elucidated, its response characteristics are still sparsely mapped.

At the molecular level, regulation of mTORC1 activity is achieved spatially by the recruitment of mTORC1 and various adaptor components to the outer surface of the lysosomal membrane (Fig. 1A)(Menon et al., 2014). When amino acids are abundant, the Ragulator-Rag complex dynamically scaffolds mTORC1 at the lysosome (Lawrence et al., 2018), where Rheb, a lysosome-associated small GTPase, activates mTORC1 catalytic activity. Growth factors exert their effect on mTORC1 through regulation of Rheb, by modulating the localization of the TSC1/2 complex, a GTPase-activating protein (GAP), at the lysosomal membrane. Phosphorylation of TSC1/2 by the growth factor-induced kinases ERK and AKT triggers its dissociation from the lysosome, permitting Rheb to activate mTORC1 (Menon et al., 2014). The energy-sensitive kinase AMPK can inhibit mTORC1 activity by both phosphorylating TSC1/2 to prevent its dissociation (Inoki et al., 2003), and by directly phosphorylating Raptor (Gwinn et al., 2008). Altogether, these details of mTORC1 activation are often interpreted to comprise a spatial “AND” gate function (Fig. 1B) that is only active when both amino acids and growth factors are present, and cellular energy status is favorable (Valvezan and Manning, 2019). This model is consistent with observations that insulin and amino acids synergistically induce phosphorylation of mTORC1 substrates measured by immunoblot, with the combination of the two stimuli achieving a level of activity that is significantly larger than the sum of the responses to each input alone (Carroll et al., 2016; Menon et al., 2014).

**Figure 1:**
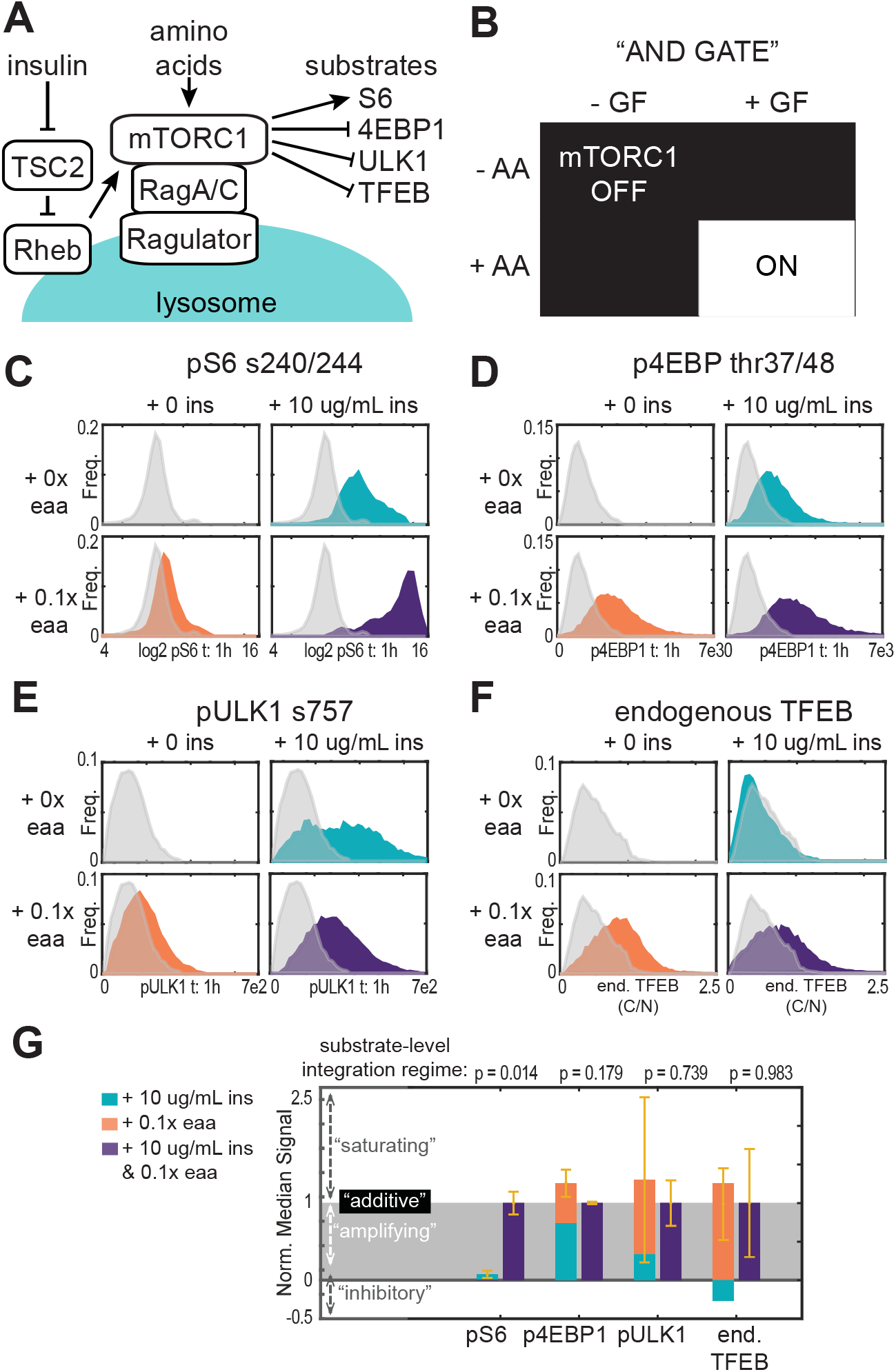
Integration of amino acids and insulin by mTORC1 substrates. A) Spatial regulation of mTORC1 activity at the lysosome. Insulin inhibits TSC2 localization at the lysosome, and thereby promotes Rheb activity. Amino acids promote Ragulator-Rag-mTOR complex formation. B) Schematic of “AND GATE” integration logic, where both insulin and amino acids are required for mTORC1 activity. (C-F) Quantification of mTORC1 target responses by immunofluorescence. MCF10A cells were starved of growth factors and amino acids for 6 hours, then stimulated for 1 hour with insulin (10 ug/mL), amino acids (0.1x), or both (vehicle control in grey). Histograms represent the distribution of single-cell values for (C) pS6 s240/244, (D) p4EBP thr37/46, (E) pULK1 s757, and (F) cytosolic to nuclear ratio of endogenous TFEB. G) Comparison of responses to individual or combined stimuli. The summed median responses of mTORC1 targets to single stimuli (red and blue stacked bars) are shown next to the observed response to combined amino acid and insulin stimulation, as measured in (C-F). Signals are normalized to the median combined response, so that a perfectly additive response to single stimuli would correspond to a value of 1. Values below 1 indicate that the observed combined response exceeds the sum of the individual responses. Error bars report variance between replicates. P values are calculated using two-tailed T-tests.

However, the molecular model of mTORC1 activation does not specify its response dynamics at the cellular level. mTORC1 and its regulators are present in >10^3^ copies per cell and can cycle rapidly between multiple states within any individual cell (Lawrence et al., 2018), potentially allowing the aggregate cellular activity of mTORC1 to vary continuously. Therefore, even if it behaves as a logical AND gate at the molecular level, mTORC1 activity at the cellular level could respond to inputs as a sharp logical switch, as a graded rheostat, or with a more complex logic. Multiple models have been proposed, but existing data are insufficient to distinguish them (Valvezan and Manning, 2019). Three factors limit the ability of previous studies to capture the activity of mTORC1 as it operates in cells. First, population averaging methods (such as immunoblots) can obscure the heterogeneous activity of single cells (Birtwistle et al., 2012; Purvis and Lahav, 2013), and most studies to date have measured mTORC1 activity by such assays rather than at the single-cell level. In an extreme example, the average mTORC1 response to two different inputs could arise from one subset of cells being activated by one input, while different cells are activated by the other, with no cell responding to both stimuli. More complex possibilities also exist, making single-cell resolution necessary. Second, mTORC1 activity is usually quantified by measuring phosphorylation of its substrates, including S6K (or its substrate, ribosomal protein S6), 4E-BP1, or ULK1 using phospho-specific antibodies. While such measurements are correlated with mTORC1 activity, this approach is limited because single time-point measurements of phosphorylation may not be linearly proportional to mTORC1 activity, and these substrates may differ in their sensitivity to mTORC1. Third, in previous studies, different stimuli are typically applied simultaneously, but this form of stimulation represents only a subset of the possible contexts in which a cell may receive stimuli. Factors such as the time between the presentation of stimuli may have an important effect on the mTORC1 response. Ideally, time-resolved measurements in individual cells could address these three limitations and more precisely characterize the cellular mTORC1 response to multiple stimuli.

A potential strategy to quantify mTORC1 activity in live cells involves the transcription factor TFEB, a member of the microphthalmia (MiTF-TFE) family of transcription factors (Goding and Arnheiter, 2019). TFEB modulates the expression of genes required for lysosomal and mitochondrial biogenesis (Mansueto et al., 2017; Settembre et al., 2013). Like many transcription factors, TFEB is regulated at the level of sub-cellular localization. mTORC1 phosphorylates TFEB at serines 122 and 211; phosphorylation at Ser-211 promotes binding to 14-3-3 proteins and sequestration of TFEB in the cytosol where it cannot access DNA (Roczniak-Ferguson et al., 2012; Settembre et al., 2012). When not phosphorylated at these serines, TFEB accumulates in the nucleus, where it activates the transcription of its target genes (Sardiello et al., 2009). The cytosolic to nuclear ratio of TFEB (hereafter TFEB_C/N_) has been frequently used as a surrogate measure of this phosphorylation, both by immunofluorescence (Marin Zapata et al., 2016) and by live-cell microscopy using a fluorescent protein (FP) fusion tag (Li et al., 2018). However, TFEB_C/N_ is also regulated by phosphorylation at additional sites. GSK3β is a potent regulator of TFEB_C/N_, phosphorylating Ser-134 and Ser-138 to enhance nuclear export (Li et al., 2018; Li et al., 2016). ERK and AKT have also been reported to phosphorylate TFEB (Palmieri et al., 2017; Settembre et al., 2012), though their impact on translocation remains more ambiguous. Using TFEB_C/N_ as a live-cell readout for mTORC1 activity depends on disentangling these complicating factors.

To address the controller function of mTORC1, it is critical to quantitatively describe its sensorial function - the mapping of input stimuli strength to output activity. Describing such functions for signaling pathways remains a general challenge (Antebi et al., 2017; Brent, 2009). What mathematical expressions are best able to represent the activity of mTORC1 as a function of multiple factors? Without more detailed measurements of these factors and activities, this remains uncertain. However, a straightforward approach is a generalized linear model (GLM), where contributions from different factors are simply added (i.e. *Y* = *a* ∗ *X*_1_ + *b* ∗ *X*_2_). If this model does not fit the available data well, “higher order” interactions can also be included, such as amplification when both factors are present simultaneously (e.g. *Y* = *a* ∗ *X*_1_ + *b* ∗ *X*_2_ + *c* ∗ *X*_1_ ∗ *X*_2_). Distinguishing these interactions requires carefully controlled quantitative data, and the effects of dynamics must be considered, especially in physiological conditions.

In this study, we set out to develop a more formal conception of the sensorial function of mTORC1 at the cellular level. We measured both endogenous substrates and engineered versions of TFEB to isolate the individual and combined effect of multiple inputs on mTORC1 activity, and we explain how the observed dynamics can be cast in terms of a generalized linear model (GLM). Our data show that the response pattern to amino acids and growth factors varies between mTORC1 substrates but displays incremental responses rather than acting as a logical AND gate. We show that such incremental behavior allows mTORC1 to vary continuously in response to nutrient limited environments, providing a more precise understanding of the cellular physiology of mTORC1.

## Results

### mTORC1 substrates vary in their quantitative responses to combined inputs

For an initial evaluation of the joint effect of amino acid and growth factor stimuli on mTORC1 activity at single-cell resolution, we performed quantitative immunofluorescence for endogenous mTORC1 targets, including 4E-BP1 (phosphorylation at Thr-37/48), ULK1 (phosphorylation at Ser -757), S6 (phosphorylation at Ser -240/244 by the direct mTORC1 target S6K), and TFEB (cytosolic/nuclear ratio). For each stain, we examined mTORC1 target staining under three stimulation conditions: saturating insulin, essential amino acids (at a concentration inducing the maximal effect), or both insulin and amino acids. Over 10^4^ cells per condition were quantified to obtain well-sampled distributions (Fig. 1, C-F and Fig. S1, A-D). The measured cellular staining intensities were distributed unimodally for most of the stimulations and stains, indicating the absence of distinctly responding subpopulations. The one notable exception was pS6, which showed a broad non-normal distribution (Figure 1C) that was visible as highly variable individual cell intensities (Figure S1A).

Based on the quantified distributions, we assessed whether each of the mTORC1 targets displayed evidence of synergistic activation by comparing the median value of the individual insulin and amino acid response distributions to the median for the combined response (Figure 1G). pS6 showed a synergistic response, as the median for the combined response far exceeded the individual inputs alone. In contrast, staining for p4EBP, pULK1 and TFEB was not significantly different between the combined input and the sums of individual inputs. Each substrate presented different responses to the individual treatments, with TFEB showing the most divergent behavior. An approximately maximal response was triggered by amino acids alone, and a moderately negative response induced by insulin alone. Together, these data indicate that the apparent synergy of mTORC1 activity depends greatly on the substrate that is monitored, and that the mTORC1-S6K1-S6 phosphorylation cascade may amplify mTORC1 activity. Because many studies of mTORC1 signaling integration have relied on pS6 or pS6K as an indicator of activity, these results motivate a broader investigation of integration by mTORC1 substrates.

### Fluorescent reporters for TFEB localization enable continuous monitoring of mTORC1 activity in living cells

To obtain temporal resolution, we examined TFEB localization as an indicator of mTORC1 activity. However, the atypical response of TFEB localization to insulin (Fig. 1F, S1D) complicates its use as a general reporter for mTORC1. Moreover, it has been reported that TFEB phosphorylation occurs selectively in response to mTORC1 activity stimulated by amino acids, but not in response to growth factors (Napolitano et al., 2020). We suspected that these complexities in TFEB’s quantitative regulation could arise from a failure to account for temporal regulation, or from non-mTORC1-specific aspects of its regulation, including sites potentially phosphorylated by GSK3β and AKT (Li et al., 2016; Palmieri et al., 2017). To understand how these factors influence TFEB localization, we engineered multiple versions of TFEB fused to a fluorescent protein and expressed them stably in MCF10A cells (Figure 2A). We first compared full length TFEB fused to mVenus (FL-TFEB-TR) to a truncated version comprising residues 1-237, which lacks the DNA-binding domain and a C-terminal AKT phosphorylation site (TFEB-TR) (Fig. 2B,C). An advantage of this truncation is that it reduces the deleterious effects of over-expressing TFEB, which were evident in altered cellular morphology and low expression level of FL-TFEB-TR in stable clones (Figure 2B). We also constructed phospho-site mutants to decouple GSK3β and mTORC1 regulation of TFEB. In 3xSA-TR, serines at positions 134, 138, and 142, which are involved in GSK3β and ERK control of TFEB_C/N_ (Li et al., 2018; Li et al., 2016), were converted to alanine. In 5xSA-TR, the aforementioned sites and the mTORC1 sites Ser-122 and Ser-211 were all converted to alanine to yield a reporter that is expected to be unresponsive to both GSK3β and mTORC1. Finally, as a parallel measurement of mTORC1-dependent translation, we constructed cell lines carrying TOP-H2B-YFP-DD (Han et al 2014). This reporter uses a chemically inhibited degron to quantify the translation rate of an mRNA carrying a 5’ terminal oligo-pyrimidine (TOP) motif, which is found in many mRNAs regulated by mTORC1 activity (Thoreen et al., 2012).

**Figure 2:**
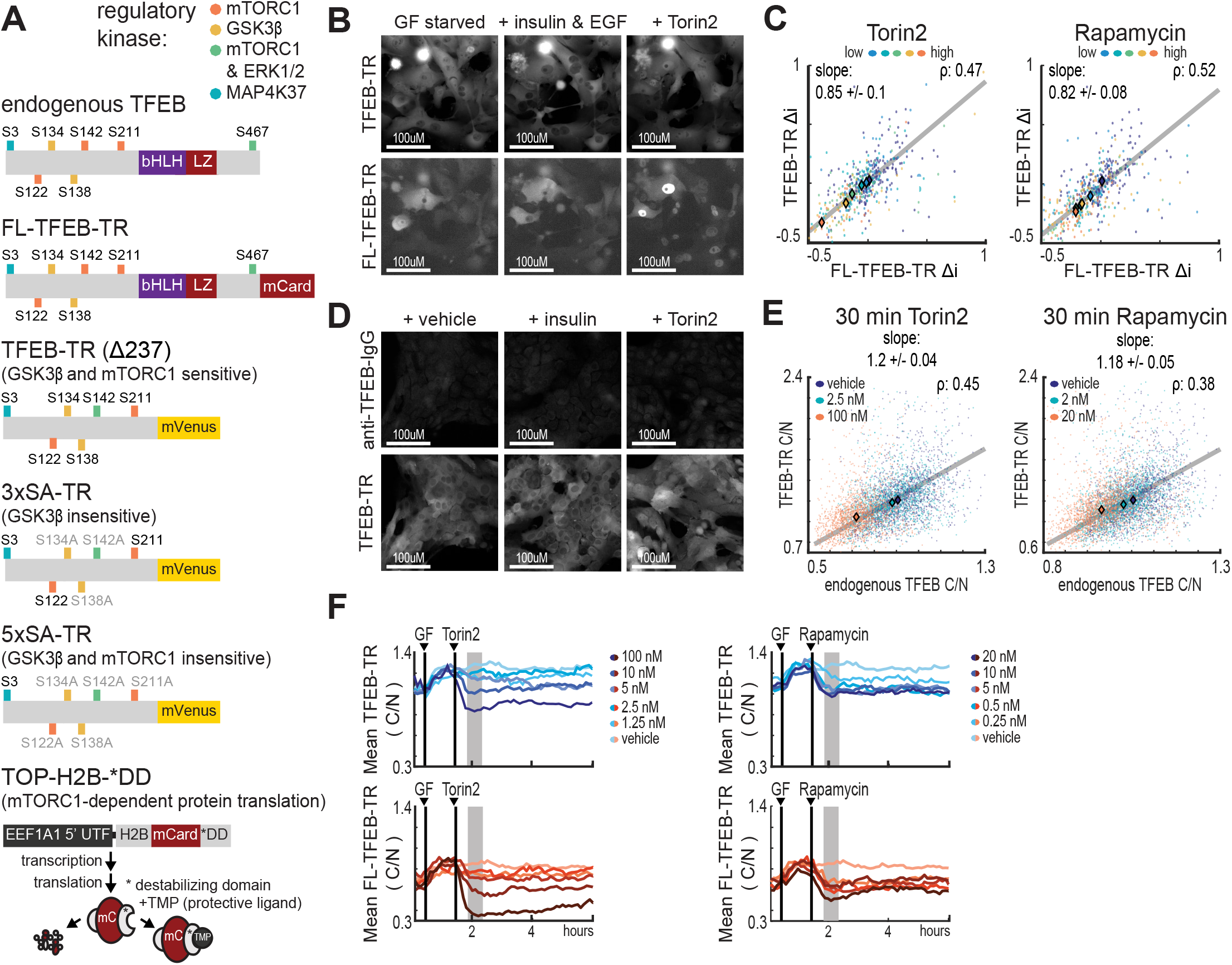
Fluorescent reporters for TFEB localization enable monitoring of mTORC1 activity in living cells. **A)** Schematic of fluorescent protein reporters used in this study. Regulatory phosphorylation sites are indicated for endogenous TFEB (top), full-length TFEB translocation reporter (FL-TFEB-TR), truncated TFEB-TR (amino acids 1-237), 3xSA-TR (lacking potential GSK3β target sites), and 5xSA-TR (lacking both GSKβ- and mTORC1 sites). The TOP-H2B-DD reporter (bottom) measures mTORC1-dependent protein translation as the rate of fluorescence increase following inhibition of degradation with TMP (Han et al., 2014). **B)** Microscopy images of MCF10A cells expressing both FL-TFEB-TR and TFEB-TR. Cells were first starved of growth factors for 6 hours, then treated for 1 hour with vehicle, EGF (20 ng/mL) and insulin (100 ng/mL), or Torin2 (100nM). Representative images from two experiments are shown. **C)** Scatter plot of same-cell Δi values for dual reporter MCF10A cells (B) treated with Torin2 or Rapamycin titrations. Δi was quantified as the mean response over a 1-hour period following treatment for >150 cells per condition. The Pearson correlation (r) for same-cell FL-TFEB-TR Δi and FL-TFEB-TR Δi TFEB-TR is indicated, along with a linear fit of the relationship and its slope. **D)** Microscopy images showing immunofluorescence detection of endogenous TFEB in TFEB-TR expressing MCF10A cells. Cells were treated as in (B) and fixed following 30 minutes of treatment. Endogenous TFEB was detected with an antibody against the C-terminal region that is absent in TFEB-TR. **E)** Scatter plot of same-cell C/N ratios for endogenous TFEB and TFEB-TR (D), representing >600 cells per condition. The Pearson correlation (r) for same-cell C/N values is reported. Linear fits of the relationships are shown with their slopes. **F)** Comparison of temporal TFEB-TR and FL-TFEB-TR localization responses to modulation. The mean C/N ratio for each reporter is shown for MCF10A cells expressing both reporters. Treatments with 20 ng/ml EGF and 10 mg/ml insulin (GF), followed by Torin2 or Rapamycin, are indicated by vertical lines. Ratios were calculated from >150 cells, with experiments performed in duplicate.

For all reporters, we quantified their responses to treatment on a cell-by-cell basis using a metric, Δi, which computes the average change in reporter localization from baseline to the peak response, typically 30-60 minutes after treatment (see Methods). Using this quantification, we observed that FL-TFEB-TR_C/N_ and TFEB-TR_C/N_ showed similar responses to a panel of mTORC1 modulators (Fig. 2B,C, S2A,B), including the inhibitors Torin-2 and rapamycin. Notably, in previous work, induction of TFEB nuclear localization by rapamycin was below the limit of detection based on either immunofluorescence or static images of TFEB-GFP (Roczniak-Ferguson et al., 2012; Settembre et al., 2012). The ability to detect responses to even low doses of rapamycin (5 nM) indicates the enhanced sensitivity that is possible using current image quantification methods. Furthermore, time-resolved data allow direct comparison of the drug-treated state to the pre-treated baseline for each cell. Similarly, the response to insulin, which was previously undetected when fixed time points were analyzed (Napolitano et al., 2020), was clearly detectable and significant using Δi as a metric (Fig. S2A). FL-TFEB-TR localization was more biased toward the nucleus than TFEB-TR, but when expressed in the same cell, both reporters showed similar dynamic changes in C/N ratio and showed high similarity by cross-correlation analysis (Fig. S2C,D,F). Based on this similarity and the reduced impact on cellular behavior, we subsequently focused on TFEB-TR.

Localization of TFEB-TR was compared to endogenous TFEB on a cell-by-cell basis using immunostaining with an antibody targeting the C-terminal region of TFEB that is absent in TFEB-TR (Figure 2D). As with FL-TFEB-TR, endogenous TFEB localized more strongly in the nucleus than TFEB-TR did, but a linear relationship was retained between the C/N ratios of TFEB-TR and endogenous TFEB over a range of kinase inhibitor treatments (Fig. 2E, S2C). Thus, TFEB-TR represents changes in the sub-cellular distribution of endogenous TFEB, without evidence of saturation. Although TFEB phosphorylation has been reported to alter TFEB protein stability (Sha et al., 2017), total TFEB-TR fluorescence changed on a slower time scale than changes in localization, suggesting that localization is the parameter most sensitive to time-dependent changes in mTORC1 activity (Fig. S2F,G). We also tested whether reporter expression alters the kinetics of mTORC1 substrate phosphorylation by measuring pS6 or p4E-BP1 in a time course, and we observed no difference in average phosphorylation kinetics between reporter-expressing and parental MCF10A cells (Fig. S2H). We conclude that TFEB-TR_C/N_ tracks with endogenous TFEB_C/N_ in live cells in response to both amino acid and growth factor stimuli, with minimal perturbation of endogenous mTORC1 signaling.

### Phosphorylation of TFEB by GSK3 amplifies cytosolic localization induced by mTOR

We next investigated the role of GSK3β in the regulation of TFEB, which involves both direct and indirect effects on TFEB localization and may exert opposing effects on TFEB downstream of AKT (Li et al., 2018; Li et al., 2016) (Fig. 3A). To assess the importance of GSK3β phosphorylation of TFEB, we compared TFEB-TR to 3xSA-TR (Fig. 3B) across a range of conditions (Fig. 3C-F). Consistent with the reported role of the mutated serines in Crm1-mediated nuclear export of TFEB (Li et al., 2018), the average C/N ratio was decreased for 3xSA-TR across all conditions. Like TFEB-TR, 3xSA-TR responded to amino acids and insulin, but did so with a reduced amplitude (Figs. 3D,F and S3C,E). C/N ratio decreased for both reporters in response to direct inhibitors of mTOR. This decrease was large relative to baseline for TFEB-TR_C/N_, whereas mTOR inhibition only returned 3xSA-TR_C/N_ to its baseline, indicating a reduction in its overall dynamic range (Fig. 3G,H). To quantify this decrease in dynamic range, we directly compared Δi for the two reporters expressed in the same cell (Fig. 3G,H, Fig. S3C,D,E). Correlation between the reporters was strong given the expected variance in single-cell measurements. However, the slopes of the relationships were considerably lower (0.21, 0.26) than the slope of 1 expected for equivalent responses. These slopes indicate a smaller change in 3xSA-TR_C/N_ than in TFEB-TR_C/N_ across all conditions. Taken together, these findings indicate that phosphorylation of TFEB by GSK3β expands the dynamic range of TFEB translocation in response to both increases and decreases in mTORC1 activity.

**Figure 3:**
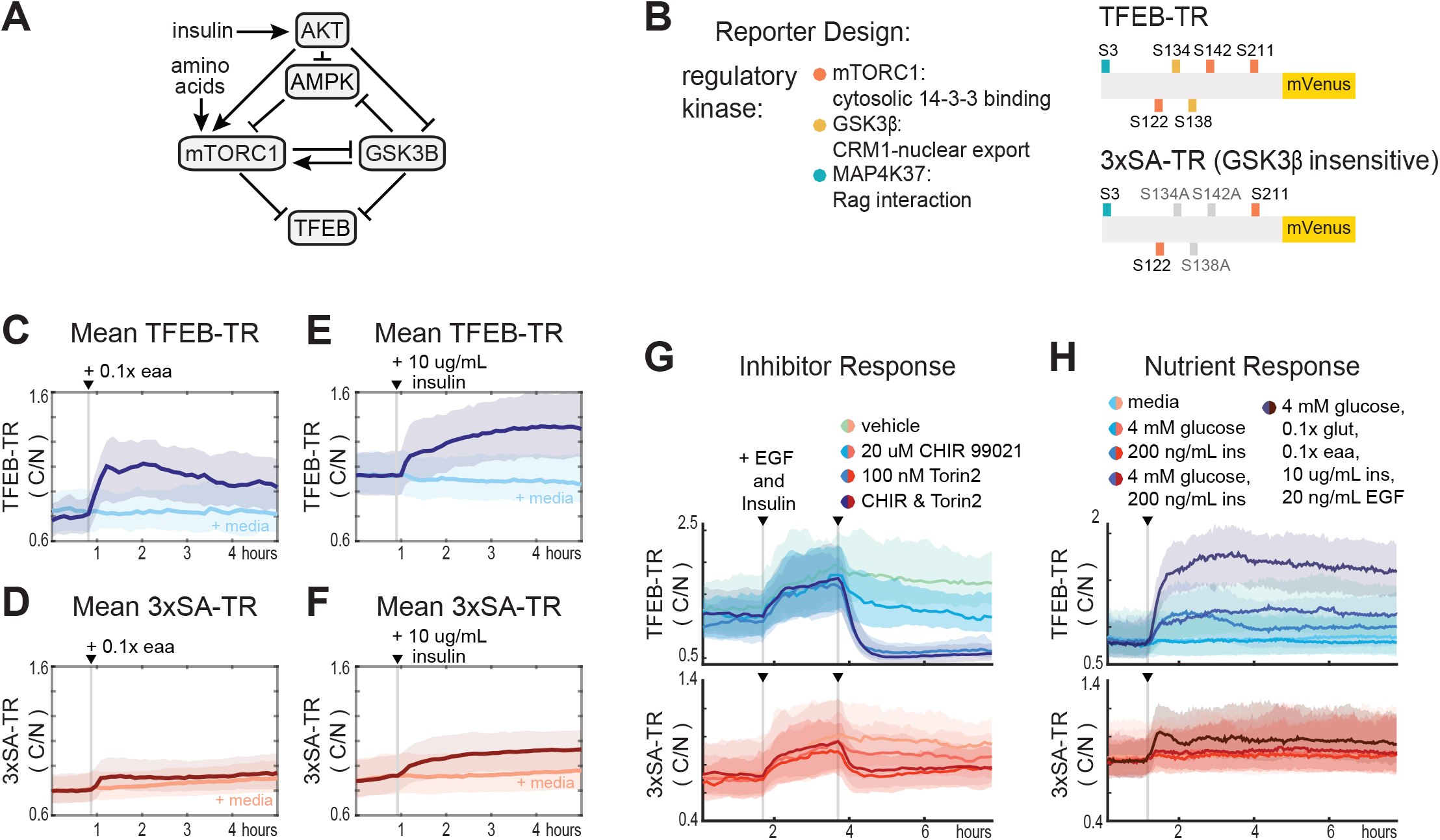
Phosphorylation by GSKβ amplifies cytosolic to nuclear ratio of TFEB. **A)** Schematic diagram of kinase interactions regulating TFEB nuclear localization. **B)** Illustration of wild type TFEB-TR, and the GSKβ-insensitive reporter, 3xSA-TR. **(C-F)** Temporal changes in reporter C/N ratio following stimulation with essential amino acids (eaa, 0.5x) or insulin (10 mg/mL). Experiments were performed in MCF10A cells expressing both TFEB-TR and 3xSA-TR. Bold lines indicate mean, with the 25th and 75th percentiles indicated by shaded regions. **(G-H)** Comparison of nutrient and inhibitor responses for both TFEB-TR and 3xSA-TR reporters, in dual reporter cells. Mean C/N ratio is indicated by bold lines with the 25th and 75th percentiles indicated by the shaded regions.

Surprisingly, despite the mutation of its GSK3β phosphorylation sites, 3xSA-TR showed a decrease in C/N ratio in response to a GSK3β inhibitor (CHIR99021, Fig. 3G). A possible explanation is indirect inhibition of mTORC1 activity through known regulation by GSK3β (Stretton et al., 2015), and consistent with this, CHIR99021 decreased staining for pS6 under a range of conditions (Fig. S3A). As expected, all responsiveness to insulin and amino acids was lost in the 5xSA-TR construct, in which the two remaining sites required for mTORC1-dependent responses (Ser-122 and Ser-211) were mutated (Fig. S3B).

### Incremental sensing of input strength by mTORC1-TFEB

Having characterized the TFEB-TR and 3XSA-TR reporters, we used them to ask how the mTORC1-TFEB pathway senses increasing concentrations of individual stimuli. This approach distinguishes sharp “digital” responses, which appear as bi-modal distributions at intermediate stimulus concentrations, from “analog” or graded responses, which are characterized by a gradual shift in the mean of a unimodal distribution (Tyson et al., 2003)(Fig. 4A). Distributions of individual cell responses were made by first withdrawing amino acids, and then calculating the Δi (peak change in the first hour) or ΔSS (change after 3-4 hours) for each individual cell (Fig. 4B) after amino acids were resupplied. Both metrics shifted gradually and unimodally with the concentration of amino acids, indicating that the individual cell response to amino acids is graded. Similar distributions were also obtained for 3xSA-TR and TOP-H2B-YFP-DD (Fig. 4C), and graded responses were also observed for other mTORC1 modulators, including insulin, and pharmacological inhibitors of mTORC1 and its upstream activators (Figs. 4D,E and S4A-I). Together, these results demonstrate that, in general, the response of mTORC1-TFEB signaling is incremental.

**Figure 4:**
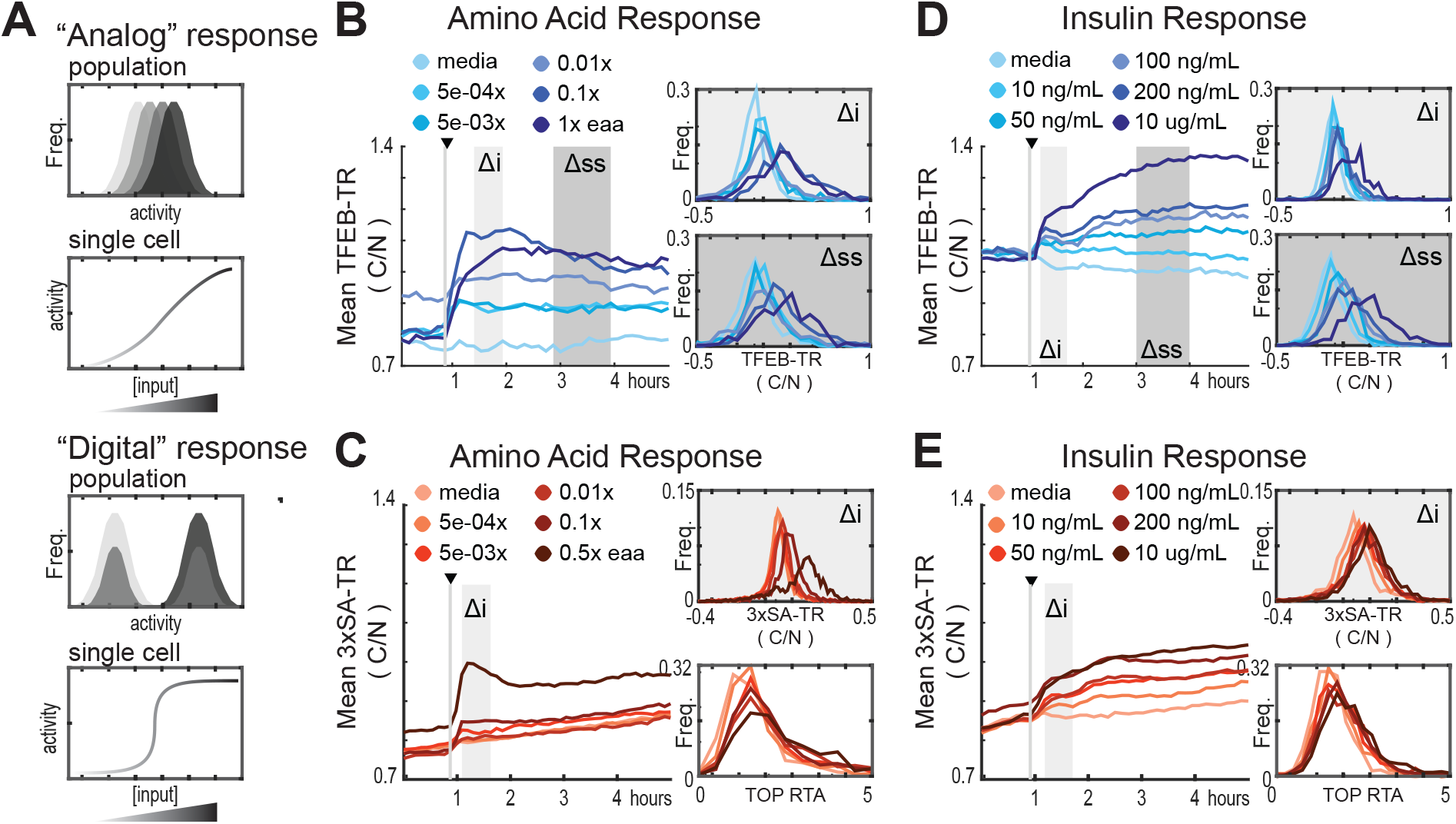
Analog regulation of mTORC1-TFEB axis by input strength. **A)** Conceptual comparison of “Analog” and “Digital” signal responses to graded input concentrations. Histograms and single-cell plots represent the expected results if individual cells show a dose-responsive (analog) behavior or a sharply switching (digital) behavior at a threshold concentration. **(B-E)** Response of MCF10A-TFEB-TR, 3xSA-TR, or H2B-TOP-DD cells to titrations of mTORC1 modulators. The mean C/N ratio over time for over >500 cells is reported as a vline, along with histograms of the initial (Δi) and steady state (Δss) responses for individual cells. (B&C) Cells were starved of growth factors and amino acids for 6 hours then stimulated with essential amino acids. (D&E) Cells were starved of growth factors then treated with titrations of insulin.

To evaluate how multiple inputs interact in the mTORC1-TFEB pathway, we took advantage of the fact that with a live-cell reporter, stimulus responses can be measured sequentially in the same cell. We developed a protocol in which cells were first cultured in the absence of sugars, amino acids, and growth factors, which were then replaced in various sequences at 2-hour intervals (Fig. 5). As internal controls for the activity of upstream mTORC1 regulators, we included reporters for AKT and AMPK (Hung et al., 2017). When glucose and glutamine were added first (Fig. 5A), an immediate increase in mean TFEB-TR_C/N_ was observed, coinciding with a decrease in mean AMPK activity. Subsequent treatment with insulin resulted in a further increase in mean TFEB-TR_C/N_, along with a corresponding activation of AKT. These responses align with basic expectations for mTORC1 regulation by TSC1/2 and Rheb. A third treatment, with amino acids, spurred another increase in mean TFEB-TR_C/N_, with no further increase in AMPK activity, and a small decrease in AKT activity, which is consistent with amino acids activating mTORC1 through a Rag-dependent mechanism independent of both AKT and AMPK.

**Figure 5:**
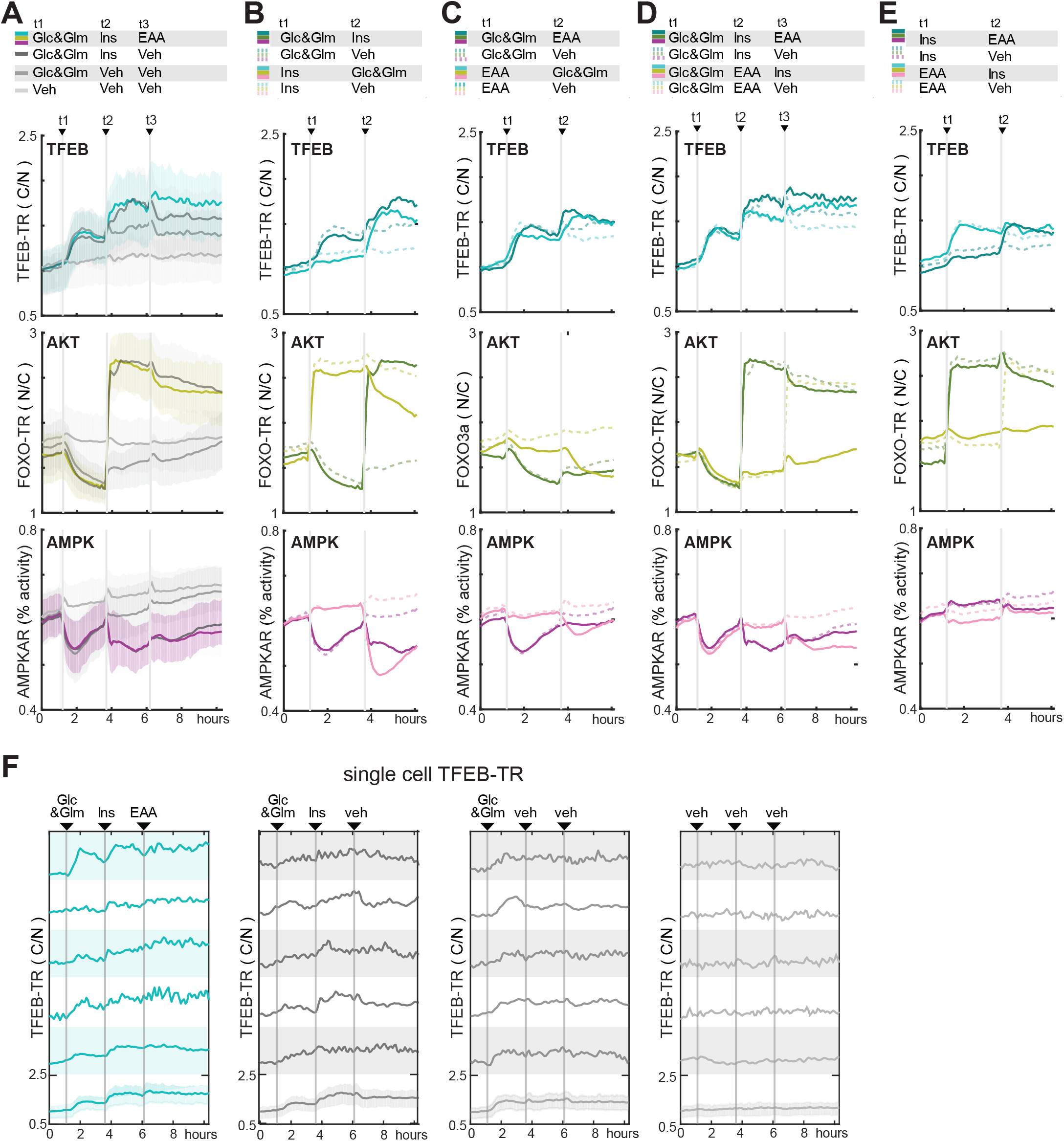
TFEB responds to sequential nutrient addition through incremental changes in localization. **A)** Mean reporter responses to sequential nutrient stimulation in MCF10A cells expressing TFEB-TR, an AKT reporter (FOXO-TR), and an AMPK activity FRET reporter (AMPKAR2). Cells were starved of nutrients and growth factors for 6 hours, then stimulated with glucose (17.5mM) and glutamine (2.5 mM), followed by essential amino acids (0.1x), and then insulin (10 mg/mL). The mean and 25th/75th percentile were calculated from >200 cells. The grey lines represent either media-only control stimuli, or glucose and glutamine, essential amino acids, then media. **B-E)** Mean reporter responses to sequential nutrient stimulation in the MCF10A triple reporter cell line. Cells were starved of nutrients for 6 hours, then stimulated with glucose (17.5mM) and glutamine (2.5 mM), insulin (100 mg/mL), and essential amino acids (0.1x) in the indicated sequences. Lines represent means of >200 cells; dotted lines show vehicle controls. **F)** Single cell responses of TFEB-TR C/N for the stimulus sequence shown in (B). Representative cells were chosen randomly, excluding outlier examples.

Individual cells also displayed incremental, stepwise changes in TFEB-TR_C/N_ in response to the sequential stimuli (Fig. 5F). However, responses to different stimuli varied in magnitude, and in many cases were difficult to distinguish from the basal variability of TFEB-TR_C/N._ These data therefore confirm that individual cells can respond incrementally to multiple mTORC1 stimuli, but also demonstrate variation in the sensitivity to amino acids, glucose and glutamine, or insulin, which may represent underlying heterogeneity in the metabolism of individual cells.

When the sequence of stimuli was modified, we noted changes in the average size of TFEB-TR responses. Notably, when insulin was added before amino acids, glucose, or glutamine, there was a minimal response of TFEB-TR_C/N_, despite a robust activation of AKT (Fig. 5B). When glucose and glutamine were subsequently introduced, the resulting TFEB-TR_C/N_ response reached a magnitude similar to that achieved by insulin following glucose and glutamine. This dependency may indicate that direct activation of mTORC1 by AKT-mediated phosphorylation is substantially weaker than by the AKT-stimulated uptake of glucose, which could directly activate mTORC1 and/or suppress AMPK activity. The activity profile for AMPK is consistent with this latter possibility. In contrast to this dependency, a substantial TFEB-TR response to amino acids was observed even prior to glucose/glutamine addition (Fig. 5C), indicating that Rag-mediated activation of mTORC1 is less reliant on extracellular glucose and glutamine. Finally, we alternated the order of amino acid and insulin stimuli in cells that had already received glucose and glutamine. TFEB-TR_C/N_ responses to insulin and amino acids were comparable regardless of their order in the sequence; we did not observe any enhancement of insulin or amino acid response by pretreatment with the other factor (Fig. 5D,E). Together, these results again diverge from a strict AND gate model and support a rheostat-like model with a continuous graded response of mTORC1. When the same experiments were repeated with the 3xSA-TR biosensor, results were consistent with those of TFEB-TR, but the diminished range of the 3xSA reporter reduced the ability to distinguish incremental shifts in reporter localization from noise (Fig. S5).

### Temporal fluctuations in mTORC1 activity track individual cell metabolic states

By monitoring cells over time, we noted that TFEB_C/N_ did not remain constant following each stimulus. In many cases, it showed a rebound effect following an initial peak or a gradual rise across hours. These transients imply that mTORC1-TFEB signaling continually adjusts to cellular metabolite availability. We therefore tracked cells over a longer period of time (12 hours) under conditions of starvation and refeeding with essential amino acids at different concentrations (Fig. 6A). Initially after refeeding, average TFEB-TR_C/N_ changed sharply depending on the amino acid concentration, as shown before. Subsequently, however, mean trajectories converged slowly approximately 12 hours after re-feeding, regardless of the amino acid concentration. Adaptation on this time scale indicates that mTORC1-TFEB signaling eventually reaches a new steady state, consistent with the expectation that TFEB-directed induction of autophagy will eventually provide more amino acids, leading to resumed activity of mTORC1.

**Figure 6:**
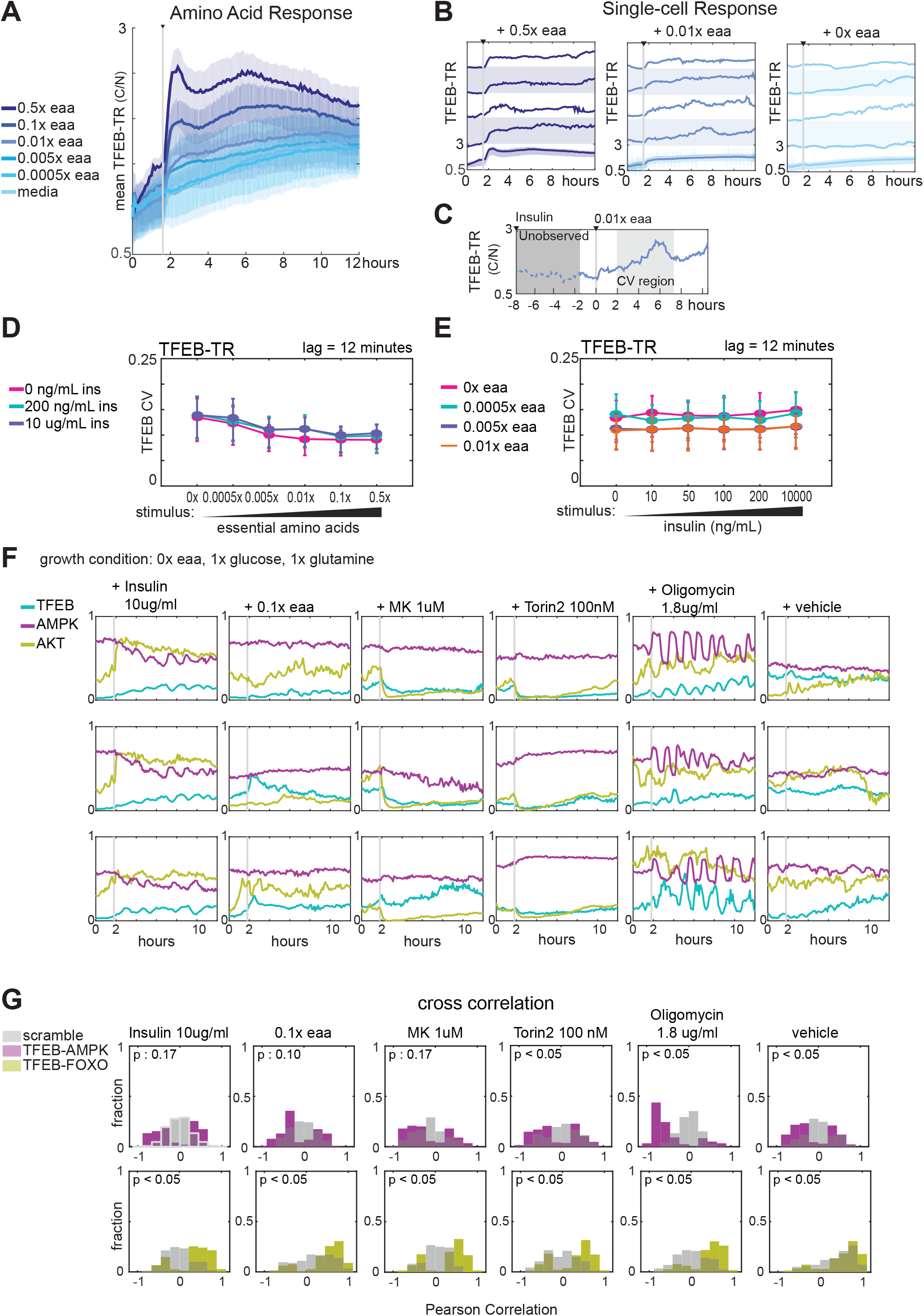
Nutrient limitation induces correlated temporal fluctuations in the AMPK-mTORC1-TFEB network. **A)** Mean TFEB-TR C/N for 1 hours in MCF10A cells, following starvation of growth factors and amino acids for 6 hours and then stimulated with essential amino acids at a range of concentrations. The mean TFEB-TR C/N is shown over 15 hours for >400 cells in each condition. **B)** Single cell TFEB-TR C/N responses, for conditions corresponding to the experiment in (A). Cells were starved of amino acids and growth factors for 6 hours then stimulated with media, 0.01x essential amino acids, or 0.5x essential amino acids. The mean N/C is indicated by the grey line with shaded regions denoting the 25th and 75th percentiles. **C)** Schematic illustrating coefficient of variation calculation for C and D. **(D,E)** Single cell, coefficient of variation (CV) in TFEB-TR for MCF10A cells grown in dual gradients of insulin and essential amino acids. CV was calculated for >400 individual cells per condition, by calculating the standard deviation of the time series divided by the mean of TFEB-TR C/N over 2-7 hours after stimulation. The error bars report the interquartile range of the CV. iD) Cells were cultured in amino acid gradients for 6 hours then stimulated with a gradient of insulin. E) Cells were cultured in a gradient of insulin for 6 hours then stimulated with a gradient of amino acids. **F)** Single cell traces for triple reporter MCF10A cells. Cells were starved of amino acids for 6 hours and then treated with Torin2, media, 10 ug/mL insulin, or 0.1x essential amino acids. **G,H)** Cross correlation of single cell TFEB-TR and AMPKAR activity (G) or TFEB-TR and FOXO-TR activity (H), calculated for >150 cells during a 2-10 hour period following treatment with 10ug/mL Insulin, 0.1x eaa, 1uM MK2206, 100nM Torin2, 1.8ug/mL Oligomycin, or vehicle treatment. Grey bars indicate “scrambled” controls in which reporter signals from different cells were paired randomly.

To observe this adaptation process at the cellular level, we analyzed TFEB-TR_C/N_ in individual cells under the same conditions. In media lacking amino acids, TFEB-TR_C/N_ fluctuated asynchronously, with a time scale of approximately two hours (Fig 6B). Unlike pulsatile signals observed in some other pathways (Albeck et al., 2013; Lahav et al., 2004), TFEB-TR_C/N_ fluctuations were not sharply defined. Therefore, to quantify fluctuations, we calculated the coefficient of variation for each individual trajectory, over 1-6 hours after the amino acid spike. These calculations identified a trend in which the mean variance per cell was reduced under increasing extracellular amino acid concentrations, such that cells showed the largest fluctuations in TFEB-TR_C/N_ when amino acids are limited (Fig. 6C). In contrast, fluctuations remained similar across all concentrations of insulin (Fig. 6D,E). We infer that the abundant availability of amino acids precludes the need for any intervention limiting mTORC1 activity, but under more physiological conditions (less than 0.01x) nutrient stress is more common, rendering mTORC1 more sensitive to its regulators.

To understand the sources of time-dependent variation in the mTORC1 signaling network, we monitored TFEB-TR_C/N_ alongside AKT and AMPK activities within individual cells over time. We observed synchronous fluctuations in these activities under multiple conditions (Fig. 6F). Oligomycin, which induces large pulses in AMPK activity, induced the most pronounced and apparently correlated activity changes in TFEB-TR, AMPK, and AKT. However, other conditions also produced fluctuations with visible relationships between the kinase activities. Cross-correlation analysis was performed to quantify these relationships between AKT and TFEB-TR, and between AMPK and TFEB-TR, revealing significant cellular co-variation. Together, this analysis indicates that AKT, AMPK and mTORC1 activity fluctuations are coupled at the cellular level in most contexts, with the largest fluctuations (other than those produced by inhibitors) observed when amino acids are limited. Overall, these data reinforce that mTORC1 activity is continually responsive to changes within its network of input pathways. The highly coupled nature of these signals demonstrates the need for a view of the metabolic signaling network that is both more global, in terms of the interacting factors, and more detailed, in the frequency of regulatory exchanges.

## Discussion

### An incremental model for multi-input sensing by mTORC1 at the cellular level

By tracking mTORC1-TFEB signaling in living cells, we provide evidence that cellular mTORC1 activity is fine-tuned, rather than abruptly induced by certain combinations of stimuli. Regardless of whether dose curves of amino acids or growth factors are presented individually or in combination, our data indicate that mTORC1 activity responds through incremental changes. Thus, we can rule out truly “digital” or “switch-like” mechanisms where the response is focused on a sharp threshold. Our single treatment dose curve experiments (Fig. 4B-D) support a model where mTORC1 activity responds in an analog format, increasing proportionally with the addition of either amino acids or insulin, which we interpret as activity of Rag and Rheb, *mTORC*1 = *a* ∗ *Rag*^*A*^ + *b* ∗ *Rheb*^*A*^.

This model has only additive terms, whereas a logical AND gate would have only the interaction term (*c* ∗ *Rag*^*A*^ ∗ *Rheb*^*A*^). While our findings might appear to conflict with the molecular AND gate model, this may be an artifact of experimental limitations. The difference can be reconciled by supposing that the basal levels of active Rag heterodimers and GTP-bound Rheb are low, but not zero (Fig. 7A, panel i). This is equivalent to stating that the *Rag*^*A*^ and *Rheb*^*A*^ model terms can never be truly zero; the observed additive terms (*a* ∗ *Rag*^*A*^, and *b* ∗ *Rheb*^*A*^) could then be explained by the minimum levels (*c* ∗ *Rag*^*A*^ ∗ *Rheb*^*A*^ _*MIN*_, and *c* ∗ *Rag*^*A*^_*MIN*_ ∗ *Rheb*^*A*^, respectively). Given this scenario, increasing *either* the activated Rag-complex binding sites (*Rag*^*A*^ ↑), or reducing the number of TSC2 complexes on the lysosome (*Rheb*^*A*^ ↑) would be sufficient to increase the number of active mTORC1 complexes (Fig. 7A, panels ii and iii). However, both stimuli would be needed to reach maximal activity. In other words, activation of each individual mTORC1 behaves as a molecular AND gate requiring engagement with both an active Rag and active Rheb, but the number of such productive complexes within the cell would be variable, allowing the total cellular activity of mTORC1 to range continuously as a sort of cellular rheostat.

**Figure 7:**
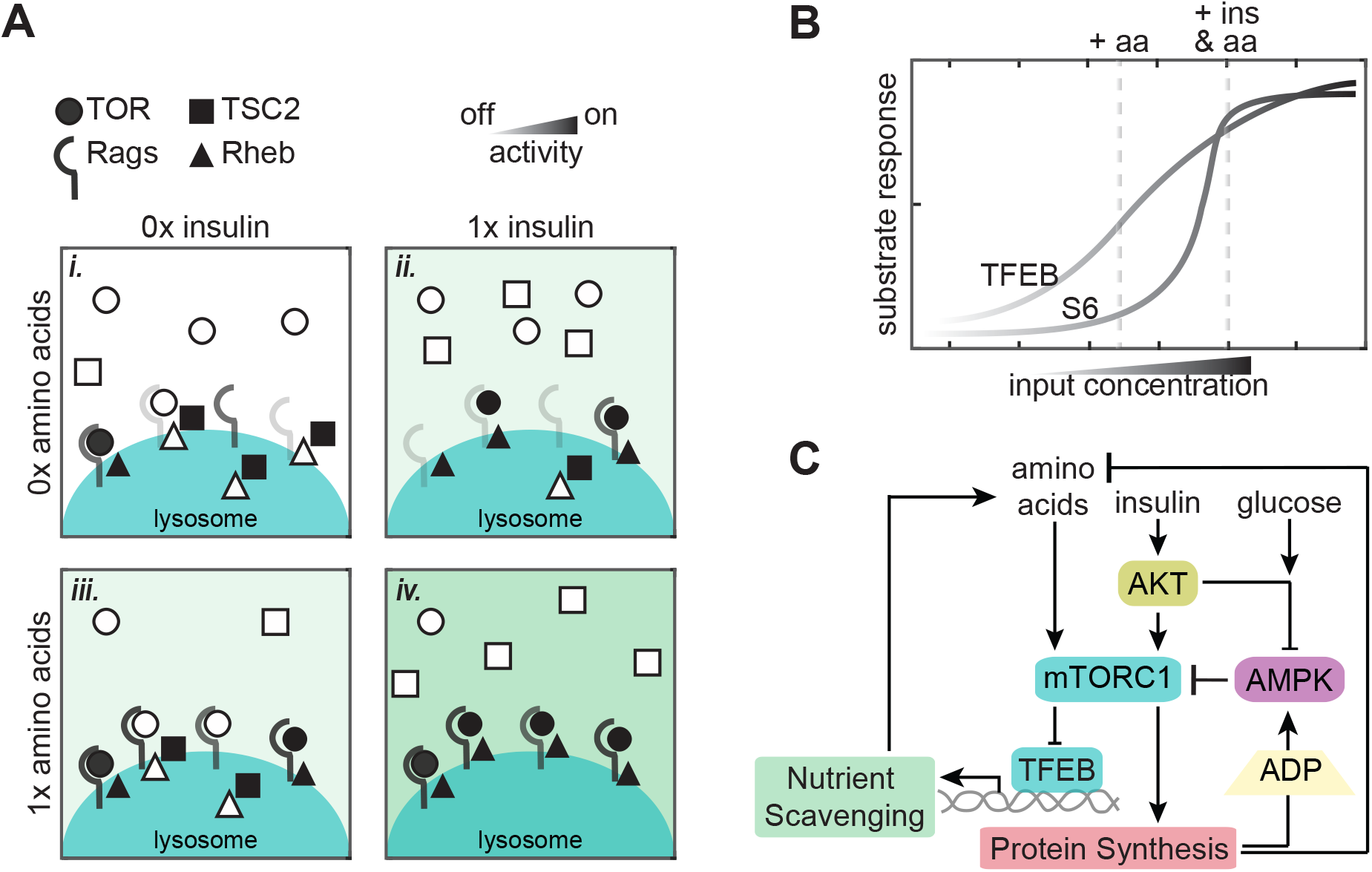
A model of continuous integration of nutrient and growth factor status by mTORC1-TFEB signaling. **A)** Schematic representation of analog integration of nutrients and growth factors by mTORC1 at the lysosome. **B)** Graph illustrating how differences in mTORC1 substrate sensitivity can change the appearance of mTORC1 integration logic. **C)** Wiring diagram illustrates potential sources for slow and fast fluctuations in mTORC1 activity. Protein synthesis produces ADP and depletes available amino acids, with potential to activate AMPK and inhibit mTORC1 when nutrients are limited. Transcriptional activity of TFEB promotes nutrient scavenging programs to balance metabolic supply and demand.

This model underscores the importance of clarifying the relationship between molecular mechanisms (i.e. how individual molecular complexes function) and systems-level behaviors (i.e. how the many copies of the complex behave collectively within the cell). The boolean logic of individual mTORC1 complexes does not imply strict on/off responses at the cellular level, but there are important cellular implications for this molecular behavior. In particular, our results using sequential stimuli (Figure 5) suggest that the spatial organization of mTORC1 activation creates a motif that prevents any single stimulus from exerting a saturating effect on mTORC1. For example, even if excess insulin saturates the insulin receptor or PI3K/AKT pathway, mTORC1 activity can remain sensitive to changes in amino acids by the independent interaction with Rag. If both growth factor and amino acid stimuli converged on the same molecular mechanism – effectively creating a molecular OR gate – it would be possible for one stimulus alone to saturate mTORC1, overriding regulation by the other stimulus. Thus, while the statistics of having many copies of each regulatory factor preclude a binary response, the molecular AND gate motif may still protect the ability of mTORC1 to remain responsive to multiple inputs.

### Differences in additive behavior between mTORC1 substrates

Our data also show that the appearance of strongly synergistic mTORC1 activation depends on which substrate is used to track its activity, and whether phosphorylation is assessed at a single time point or over time. Notably, phosphorylation of S6K or its substrate S6 have been used in most studies that have observed a strict requirement for both amino acids and growth factors (Carroll et al., 2016; Menon et al., 2014). However, we find that among the substrates we tested, pS6 is the most atypical at the single cell level, with an unusually high degree of cell-to-cell heterogeneity. A broader view of multiple substrates suggests that the activity of mTORC1 itself is graded, and that substrates vary in their non-linear response to this activity. Differences between mTORC1 substrates have been explored previously, revealing that the affinity of mTORC1 for different phosphorylation sites predicts their responses to pharmacological inhibition of mTOR (Kang et al., 2013). These differences suggest a simple model for the variation we observe between substrates, based on typical sigmoidal dose response curves (Fig. 7B). A super-additive effect would be expected for weaker substrates such as S6K, because each individual stimulus would lie within the threshold region of the response curve, and the response to the combined stimuli would be amplified by the upward concavity of the sigmoid. For more sensitive substrates, both individual stimuli and their combination would lie within the approximately linear region of the response curve, resulting in an additive effect. Based on measurements of substrate affinity, this model could explain the differences in additivity between 4E-BP1 or ULK1 and S6K1 (Kang et al., 2013).

The response of TFEB to mTORC1 is complicated by its multiple phosphorylation sites. In particular, the cluster of Ser-134, Ser-138, and Ser-142 enables glucose-sensitive regulation of TFEB localization through GSK3, mTORC2, and PKC, independently of mTORC1 (Li et al., 2018; Napolitano et al., 2018). This mechanism, which operates by regulating TFEB export from the nucleus by Crm1, in parallel to the regulation of cytoplasmic retention by 14-3-3 binding to Ser-211, could allow TFEB itself to integrate multiple inputs downstream of mTORC1. With both mechanisms operating simultaneously, it is unclear how much input integration is performed by mTORC1, rather than by TFEB. Notably, however, we observe that mutation of the GSK3β-linked sites, or withdrawal of glucose, does not abrogate the ability of TFEB to respond incrementally to multiple stimuli (Fig. S5). We infer that GSK3β-mediated phosphorylation of TFEB is not strictly essential for integrating multiple stimuli, but rather serves to amplify the responsiveness of TFEB to mTORC1. An interpretation of previous data implied that GSK3β regulation is strictly required for TFEB nuclear export (Li et al., 2018), however we argue that the two studies are reconcilable, again by interpreting all of the available data quantitatively. TFEB exchanges continuously between the nucleus and cytoplasm (Li et al., 2018; Napolitano et al., 2018), and nuclear export of Ser-142 or Ser-138 mutants is reduced partially (by 75%) rather than fully eliminated (Napolitano et al., 2018), indicating a balance between the Ser-211/14-3-3 and Ser-142/Crm-1 mechanisms rather than a strict reliance on either one. In this view, GSK3β can be thought of as an amplifier for TFEB responsiveness, which parallels the finding that GSK3β enhances the activity of mTORC1 on other substrates, including S6K1 and 4E-BP (Shin et al., 2014; Shin et al., 2011), consistent with a general role for GSK3β as a signaling amplifier of previously primed motifs (Hermida et al., 2017).

### Operational analysis of mTORC1 and implications for physiology

Physiologically, mTORC1 activity is required for embryonic development, but its inactivation is also essential. Mice deficient for TSC1 or TSC2 cannot suppress mTORC1 upon growth factor withdrawal and die during embryogenesis; *in vitro* studies of this model suggest that cell death results from a failure to limit energy expenditure (Choo et al., 2010). In mice engineered to express a GTPase-deficient (GTP-locked) RagA, mTORC1 remains active upon nutrient starvation, and the resulting death of neonatal mice is linked to a failure to induce autophagy (Efeyan et al., 2013). These studies indicate that mTORC1 inhibition during nutrient or growth factor deprivation is required to retain organismal viability during development. Under a continuous (or rheostat) model, mTORC1 activity would adapt to the abundance of the required nutrients on the time scale of both rapid metabolic fluctuations and longer-term adaptive processes. Our data, especially the correlation of TFEB-TR_C/N_ with fluctuations in AMPK activity, further support the notion that under physiological conditions, cellular energetic challenges occur continually, and over multiple time scales. For example, a period of mTORC1 inhibition may slow the rate of protein production while enabling accumulation of amino acids, induction of TFEB target genes, and a subsequent phase of mTORC1 re-activation and protein translation.

The critical function of mTORC1 is in maintaining homeostasis at the cellular level, and the present study refines the model for how it responds to several regulating factors. However, the operational behavior of the extensive feedback system in which mTORC1 operates remains relatively poorly characterized. Via the multi-reporter measurement in this study, we now conceptualize cell-intrinsic homeostasis not as a group of discrete alarm-like pathways, but as a well-connected system akin to whole-body feedback systems such as glucose or electrolyte regulation. To date, a major impediment to building on this more holistic concept has been the technical limitation on making detailed dynamic measurements at such small scales. Fortunately, a growing body of single cell techniques is beginning to enable such characterization and empower the application of quantitative models. An immediate and critical challenge to address is further multiplexing single-cell techniques to expand the repertoire of live-cell datasets that link multiple signals with metabolic changes.

## Acknowledgements

Funding for this work was provided by the National Institute of General Medical Sciences (1R01GM115650). Flow-cytometry services were supported by the UC Davis Comprehensive Cancer Center Support Grant (CCSG) awarded by the National Cancer Institute (NCI P30CA093373). We thank Roberto Zoncu for providing plasmids encoding TFEB and for helpful discussions, and Tobias Meyer for generously providing the TOP-H2B-YFP-DD reporter.

## Materials and Methods

### Immunofluorescence microscopy

For all imaging experiments, MCF10A cells were seeded in glass-bottom 96-well plates (Cellvis P96-1.5H-N, Mountain View, CA) one day prior to imaging. To increase the cell-to-media ratio, cells were plated in the center of the well by spotting 9000 cells on 3uL of type I collagen (Gibco A10483-01). Following treatment, cells were fixed in 4% paraformaldehyde solution for 15 minutes, and then permeabilized with 100% methanol for 15 minutes.

For immunostaining, cells were washed in PBS-T (0.1% Tween-20 in PBS) twice, blocked for 1 hour with Odyssey Blocking Buffer (Li-Cor, Lincoln, NE), incubated with primary antibodies overnight at 4 °C (pS6 Ser240/244 Cell Signaling Technologies (CST) #5364 dilution 1:500, p4EBP Thr37,46 CST #2855 dilution 1:200, pULK1 Ser757 CST #14202 dilution 1:800, TFEB CST #37785 dilution 1:600), and then stained with secondary antibodies (Alexa555- or Alexa647-congugated anti-rabbit, Life Technologies, A-21245, 1:250 dilution) and Hoechst-33342 (Life Technologies, H3570, diluted at 1:1000 in PBS). Images were collected with a Nikon (Tokyo, Japan) 20/0.75 NA Plan Apo objective on a Nikon Eclipse Ti inverted microscope, equipped with a Lumencor SOLA or Lumencor SPECTRA X light engine.

### Reporter construction

Plasmids encoding full-length human TFEB (FL-TFEB) were received as a gift from the Zoncu lab. The series of TFEB reporters, including FL-TFEB were constructed in the pLJM1 lentiviral vector, and the coding region of TFEB was modified, as shown in Figure 2. C-terminus of these modified TFEB constructs were fused with either mVenus or mCardinal for visualization.

TFEB-TR was constructed by truncating FL-TFEB to amino acid 1-247. 3xSA-TR was constructed by site-directed mutagenesis of amino acid residue S134A,S138A and S142A on TFEB-TR construct. Similarly 5xSA-TR, was constructed by site-directed mutagenesis of amino acid residue S122A, S134A,S138A, S142A and S211A on TFEB-TR construct.

TOP-H2B-YFP-DD was received as a gift from the Meyer lab, and is constructed and used as described in (Han et al., 2014)

FOXO-TR was constructed as described in (Hung et al., 2017) with the fluorophore changed to mOrange for spectral compatibility and expressed using retroviral transduction.

AMPKAR2 construction is as described in Hung et al., 2017 and was expressed in the pPBJ, piggyBAC transposase delivery system (Yusa et al., 2011)

### Cell line creation

Retroviral or PiggyBac systems were used to generate stably expressing reporter cell lines. Following transfection, cells were selected for using puromycin (1–2 μg/ml), G418 (200 ug/mL), or geneticin (300 μg/ml). To reduce variability in reporter expression, clonal cell lines were established using limited dilution cloning. In this study, data for each reporter is representative of behaviors that were consistent across at least three single-cell clones. Reporter cell lines were confirmed mycoplasma-negative through periodic monitoring by PCR and validation with ATCC testing of selected lines.

### Cell culture

MCF10A cells clone 5E (Janes et al., 2010) were routinely maintained in ‘DMEM/F12 growth media’ (see Media table) as described in (Debnath et al., 2003).

For live-cell time lapse microscopy experiments, we used a custom formulation, termed ‘imaging base-DMEM/F12’, which consists of DMEM/F12 lacking glucose, glutamine, riboflavin, folic acid, and phenol red (Life Technologies or UC Davis Veterinary Medicine Biological Media Service) to avoid fluorescence background. All experiments that did not involve amino acid perturbation were performed in ‘Imaging medium 1’ (see Media table). ‘Imaging medium – noAA’ was used in experiments that involves amino acid perturbation (see Media table).

Before imaging, cells were washed twice with their respective media and then cultured in imaging experiment media at least 2 hours prior to imaging. Experiments that involved amino acid perturbation, cells were pre-incubated with ‘Imaging medium – noAA’ for at least 6 prior to imaging to ascertain that cells have used all of extra amino acid from routine cell culture media.

### \Live-cell fluorescence microscopy

For live-cell microscopy experiments, cells were cultured and seeded as described in previous sections. Time-lapse wide-field microscopy was performed as detailed in Pargett et al., 2017, with cells maintained in 95% air and 5% CO2 at 37 in an environmental chamber. Images were collected with a Nikon (Tokyo, Japan) 20/0.75 NA Plan Apo objective on a Nikon Eclipse Ti inverted microscope, equipped with a Lumencor SOLA or Lumencor SPECTRA X light engine. Fluorescence filters used in the experiment are: DAPI (custom ET395/25x - ET460/50m - T425lpxr, Chroma), CFP (49001, Chroma), YFP (49003, Chroma), Cherry (41043, Chroma) and Cy5 (49006, Chroma). Images were acquired using Andor Zyla 5.5 scMOS camera every 6 – 7 minutes with at 2×2 binning. Exposure times for each channel were 25-50 ms for DAPI; 150 – 250 ms for CFP; 150 – 250 ms for YFP; 300 – 500 ms for Cherry and 300 – 500 ms for Cy5.

### Image processing and times-series data selection criteria

Live-cell microscopy images were processed using the custom MATLAB procedure as described in our previous work (Pargett et al., 2017). In this procedure, the nuclear and cytoplasmic region of individual cells are segmented, and the fluorescent intensity of pixels within the masked regions are averaged. For this study, the segmentation protocol was modified to restrict the cytoplasmic masks to the region within 5 pixels from the nuclear border, and the nuclear masks were eroded 2 pixels away from the cytoplasmic border. For all imaging experiments shown, a minimum of 100 cells were imaged and tracked for each condition.

The single-cell time series data were further filtered using minimum fluorescent intensity more than 2 times background intensity value, trace length at least 8 hours, and each trace must have no more than 3 contiguous missing datapoints. Missing datapoints in time series were linearly extrapolated to fill the gap.

### Reporter activity analysis

For all translocation reporters (FL-TFEB, TFEB-TR, 3xSA-TR, 5xSA-TR and FOXO-TR reporter), the activity of the kinases are described by the ratio of average cytosolic fluorescent intensity divided by average nuclear fluorescent intensity.

For the AMPKAR2 reporter, AMPKAR2 phosphorylation status was calculated using the protocol described in (Kosaisawe et al., 2021). Briefly, linearized AMPKAR2 FRET efficiency was calculated as shown in our previous work (Gillies et al., 2020). Then AMPKAR2 phosphorylation status was estimated based on this equation

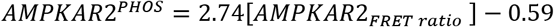

### Time series volatility analysis

For all volatility analysis of time series data, we choose time window between 2-6 hours post final-perturbation. We assume that in this time-window the kinase activity of interest has approach the quasi-steady state. Within that time window, data from every other time step were then used to calculate coefficient of variance (CV), which we used as metric of volatility. The reason that we use only data from every other time-step rather than all time-step is to avoid non-biological noise, such as imaging and segmentation noise.

### Cell trace examples

The displayed time series were chosen by random number generation in MATLAB with a threshold for minimum tracking time to eliminate cells in which recording was terminated prematurely due to failure of the tracking algorithm. The chosen tracks were manually verified to be representative of successfully tracked cells and consistent with the overall range of cell behaviors. Cell recordings determined by manual inspection to have poor tracking or quantification accuracy were discarded.

### Linear fitting and statistical tests

Linear regression analysis was preformed using a MATLAB implementation of the Theil-Sen estimator, which is insensitive to outliers and robust to heteroskedastic data.

Unless otherwise indicated, each statistical comparison was made by *t*-test with unequal variances. p-value of 0.05 is considered as significant for all hypothesis testing.

### Inhibitors

| Name | Solvent | Vendor | Catalog number |
| --- | --- | --- | --- |
| Torin2 | DMSO | Selleckchem | S2817 |
| Rapamycin | DMSO | Selleckchem | S1039 |
| Essential amino acid | Water | Gibco | 11130051 |
| MK2206 | DMSO | Selleckchem | S1078 |
| CHIR99021 | DMSO | Tocris | 4423 |
| A769662 | DMSO | Tocris | 3336 |
| Glucose | Water | Fisher Scientific | D16-500 |
| L-Glutamine |  | Thermo Fisher | A2916801 |
| Oligomycin A | DMSO | Sigma | 75351 |

### Cell Lines

| Number | Name | Source | Identifier |
| --- | --- | --- | --- |
| 1 | MCF10A | (Hung et al, 2017) | RRID:CVCL_0598 |
| 2 | MCF10A_FL-TFEB-TR | This report |  |
| 3 | MCF10A_TFEB-TR | This report |  |
| 4 | MCF10A_3XSA-TFEB-TR | This report |  |
| 5 | MCF10A_5XSA-TFEB-TR | This report |  |
| 6 | MCF10A_TFEB-TR_FOXO-TR_AMPKAR2 | This report |  |

### Media composition

*Media composition* **DMEM/F12 growth medium**
| Component | Stock Concentration | Volume | Unit | Vendor | Catalog number | Final Concentration |
| --- | --- | --- | --- | --- | --- | --- |
| DMEM/F2 | - | 500 | ml | Gibco | 11320-033 | - |
| Horse Serum | - | 25 | ml | Invitrogen | 16050-122 | 5% |
| EGF | 100 ug/ml | 100 | ul | Peptotech | AF-100-15 | 20 ng/ml |
| Hydrocortisone | 1 mg/ml | 250 | ul | Sigma | H0888 | 0.5 mg/ml |
| Cholera toxin | 1 mg/ml | 50 | ul | Sigma | C8052 | 100 ng/ml |
| Insulin | 10 mg/ml | 500 | ul | Sigma | I9278 | 10 ug/ml |

**Imaging medium**
| Component | Stock Concentration | Volume | Unit | Vendor | Catalog number | Final Concentration |
| --- | --- | --- | --- | --- | --- | --- |
| Imaging base-DMEM/F12 | - | 500 | ml | Gibco or UC Davis Veterinary Medicine Biological Media Service | Custom; equivalent to Gibco 11320-033 lacking glucose, glutamine, pyruvate, riboflavin, folic acid, and phenol red | - |
| BSA | 2% w/v | 25 | ml | Invitrogen | 16050-122 | 0.1% w/v |
| Hydrocortisone | 1 mg/ml | 250 | ul | Sigma | H0888 | 0.5 mg/ml |
| Cholera toxin | 1 mg/ml | 50 | ul | Sigma | C8052 | 100 ng/ml |
| Penicillin-Streptomycin | 10000 U/ml | 5 | ml | Gibco | 15140122 | 100 U/ml |
| L-Glutamine | 200 mM | 6.25 | ml | Gibco | A2916801 | 2.5 mM |
| Folic acid | 10 mg/ml | 212 | ul | Sigma | F8758 | 4.24 ug/ml |
| Riboflavin | 10 mg/ml | 16.25 | ul | Sigma | R9504 | 0.352 ug/ml |

**Imaging medium - noAA**
| Component | Stock Concentration | Volume | Unit | Vendor | Catalog number | Final Concentration |
| --- | --- | --- | --- | --- | --- | --- |
| Imaging base-DMEM/F12-noAA | - | 500 | ml | Gibco | Custom; equivalent to Gibco 11320-033 lacking glucose, glutamine, amino acids, pyruvate, riboflavin, folic acid, and phenol red | - |
| BSA | 2% w/v | 25 | ml | Invitrogen | 16050-122 | 0.1% w/v |
| Hydrocortisone | 1 mg/ml | 250 | ul | Sigma | H0888 | 0.5 mg/ml |
| Cholera toxin | 1 mg/ml | 50 | ul | Sigma | C8052 | 100 ng/ml |
| Penicillin-Streptomycin | 10000 U/ml | 5 | ml | Gibco | 15140122 | 100 U/ml |
| L-Glutamine | 200 mM | 6.25 | ml | Gibco | A2916801 | 2.5 mM |
| Folic acid | 10 mg/ml | 212 | ul | Sigma | F8758 | 4.24 ug/ml |
| Riboflavin | 10 mg/ml | 16.25 | ul | Sigma | R9504 | 0.352 ug/ml |

---

**Figure S1:**
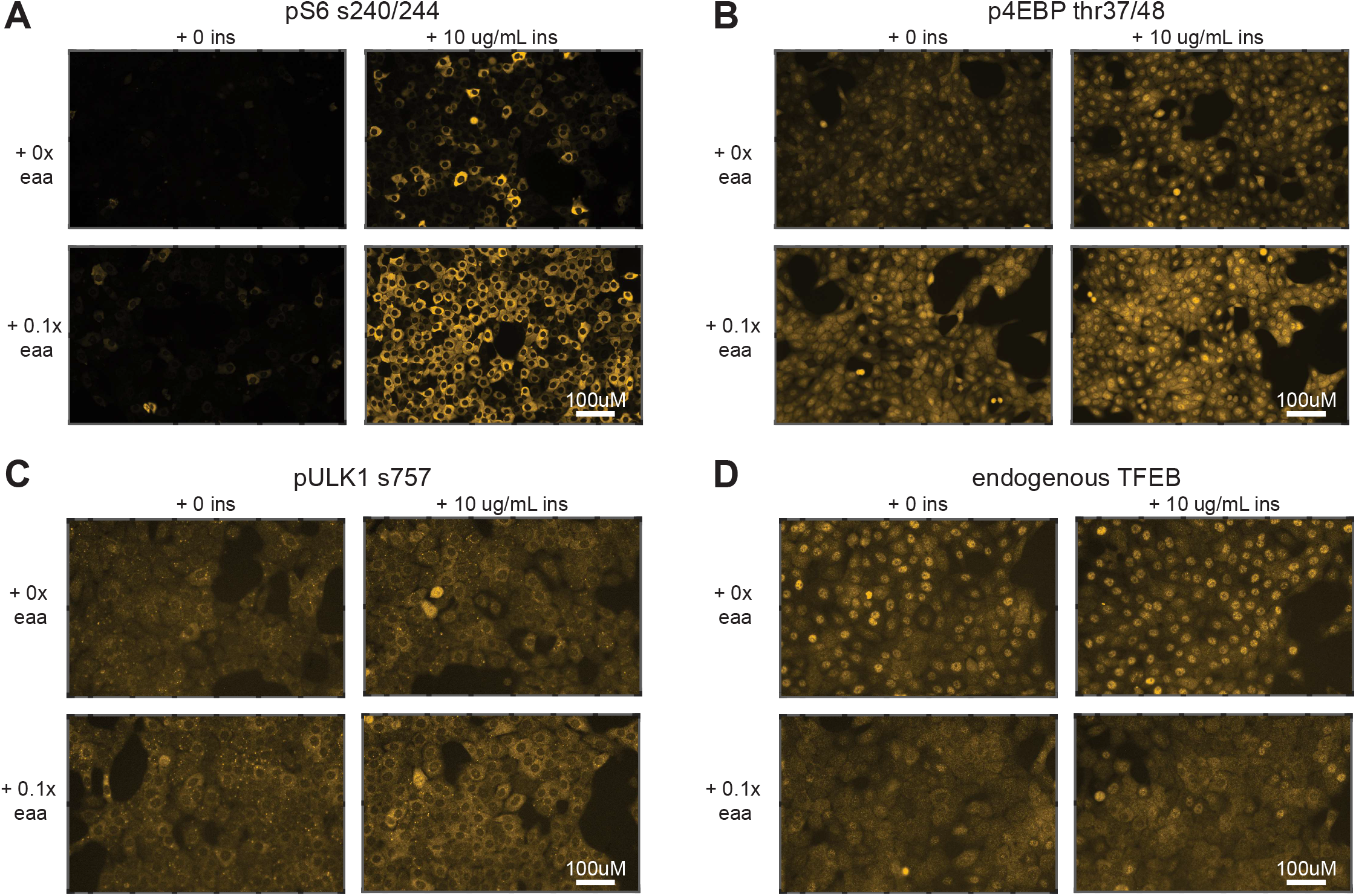
**A-D)** Immunofluorescence images of mTORC1 targets, corresponding to the quantified values in Fig. 1. MCF10A cells were starved of amino acids and growth factors for 6 hours, then stimulated for 1 hour with insulin, amino acids, or both. Images are representative examples taken from a cropped sub-region of duplicate wells.

**Figure S2:**
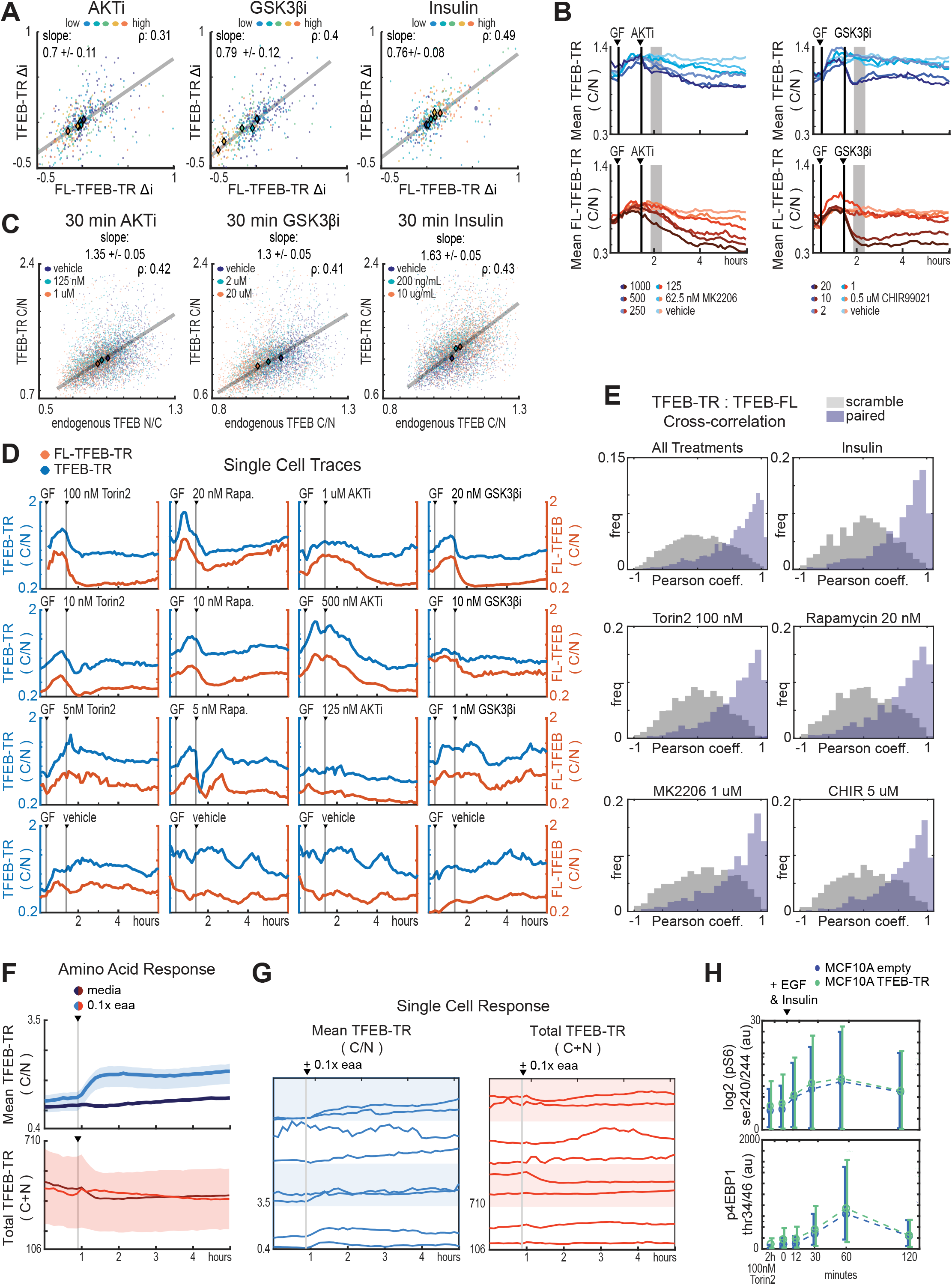
**A)** Scatter of Δi values for FL-TFEB-TR and TFEB-TR in dual reporter MCF10A cells treated with AKT inhibitor (MK2206), GSKβ inhibitor (CHIR9902) or insulin titrations. Pearson correlations and linear fits are reported as in Fig. 2C. **B)** Comparison of TFEB-TR and FL-TFEB-TR localization (mean C/N ratio) over time for dual reporter MCF10A cells treated with 20 ng/mL EGF and 100 ng/mL insulin, followed by AKT inhibitor (MK2206) or GSKβ inhibitor (CHIR9902). Ratios were calculated from >150 cells. **C)** Scatter plots of C/N ratios for endogenous TFEB immunostaining and TFEB-TR in MCF10A cells. >600 cells per condition were fixed 30 minutes after treatment with AKT inhibitor (MK2206), GSKβ inhibitor (CHIR99021) or insulin at the concentrations indicated. Pearson correlations and linear fits are reported as in Fig. 2E. **D)** Single cell traces for TFEB-TR and FL-TFEB-TR C/N in cells expressing both reporters and treated with the indicated conditions. Representative cells were chosen randomly, excluding outlier specimens. **E)** Cross correlation of single cell TFEB-TR and FL-TFEB-TR C/N ratios, calculated for >150 cells during a 5 hour period following treatment with 100nM Torin2, 20nM Rapamycin, 1mM MK2206, or CHIR99021. Grey bars indicate “scrambled” controls in which reporter signals from different cells were paired randomly. **F)** Comparison of TFEB-TR C/N ratio and total TFEB-TR fluorescence (C+N) over time. MCF10A cells were starved of amino acids for 6 hours then treated with essential amino acids (0.1x). Means are indicated by lines, with shaded areas indicating 75th and 25th quantiles. **G)** Single-cell traces of TFEB-TR C/N and total TFEB-TR fluorescence, for cells treated as in (F). Representative cells were chosen randomly, excluding outlier examples. **H)** Comparison of pS6 and p4EBP kinetics in MCF10A reporter-expressing and non-expressing cell lines. Cells were starved of growth factors for 48 hours then stimulated with EGF (20 ng/mL) and insulin (10ug/mL) prior to fixation at the indicated times. Error bars indicate standard deviation.

**Figure S3:**
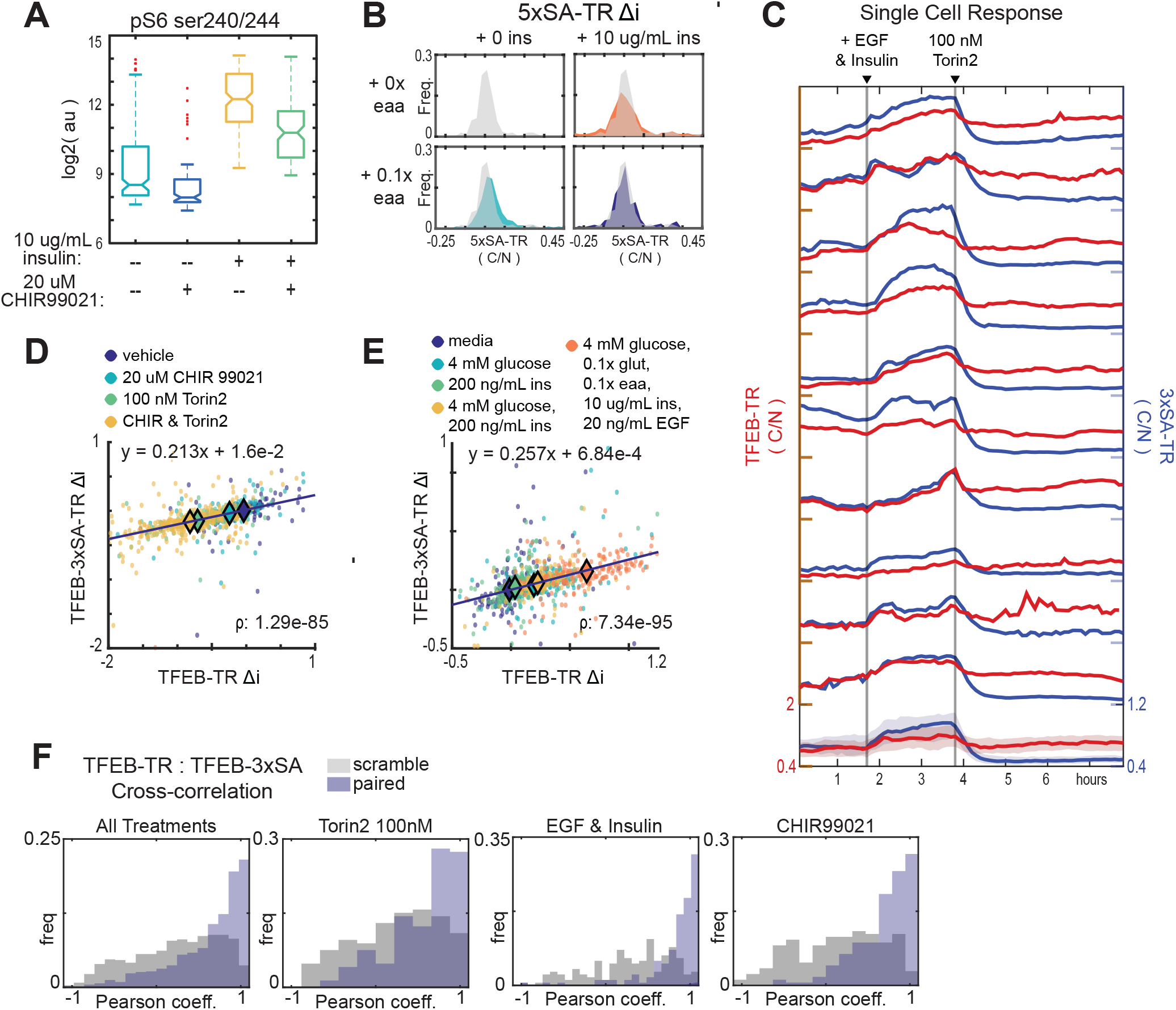
**A)** Quantification of pS6 s240/244 immunofluorescence in MCF10A cells treated with the GSK3β inhibitor, CHIR 99021, in the presence or absence of insulin. Note that pS6 intensity is decreased by GSK3β inhibition regardless of the presence of insulin, suggesting that the inhibitor could affect mTORC1 activity directly. **B)** Quantification of C/N ratios for 5xSA-TR in response to insulin or amino acid stimulation. Histograms represent the distribution of individual cell C/N ratios; the unstimulated distribution is shown as an overlay in each plot for comparison. **C)** Comparison of same-cell traces of TFEB-TR and 3xSA-TR C/N ratios, in dual reporter cells. As in Figure 3G, cells were first stimulated with EGF (20 ng/mL) or insulin (10 mg/mL), then treated with (Torin2 100 nM). **D-E)** Scatter plots of Δi for dual reporter cells. The Pearson correlation for same-cell Δi TFEB-TR and Δi 3xSA-TR is reported, and linear fits are shown with their slopes. The observed slopes of <1 indicate an attenuated dynamic range of the 3xSA-TR reporter relative to TFEB-TR. (D) Inhibitor combinations, including vehicle, GSKβ inhibitor (CHIR99021 20 mM), mTOR inhibitor (Torin2 100 nM), or both inhibitors. (E) Stimulus combinations, following 6 hour nutrient and growth factor starvation, including glucose (4 mM), insulin (200 ng/mL), both, or maximal activation by a combination of all nutrients (glucose (4 mM), glutamine (0.25 mM), essential amino acids (0.1x), insulin (10 mg/mL) and EGF (20 ng/mL). **F)** Cross correlation of single cell TFEB-TR and 3xSA-TR ratios calculated for >150 cells during a 5 hour period following treatment with 100 nM Torin2, 20 nM Rapamycin, 1 mM MK2206, or CHIR99021. Grey bars indicate “scrambled” controls in which reporter signals from different cells were paired randomly.

**Figure S4:**
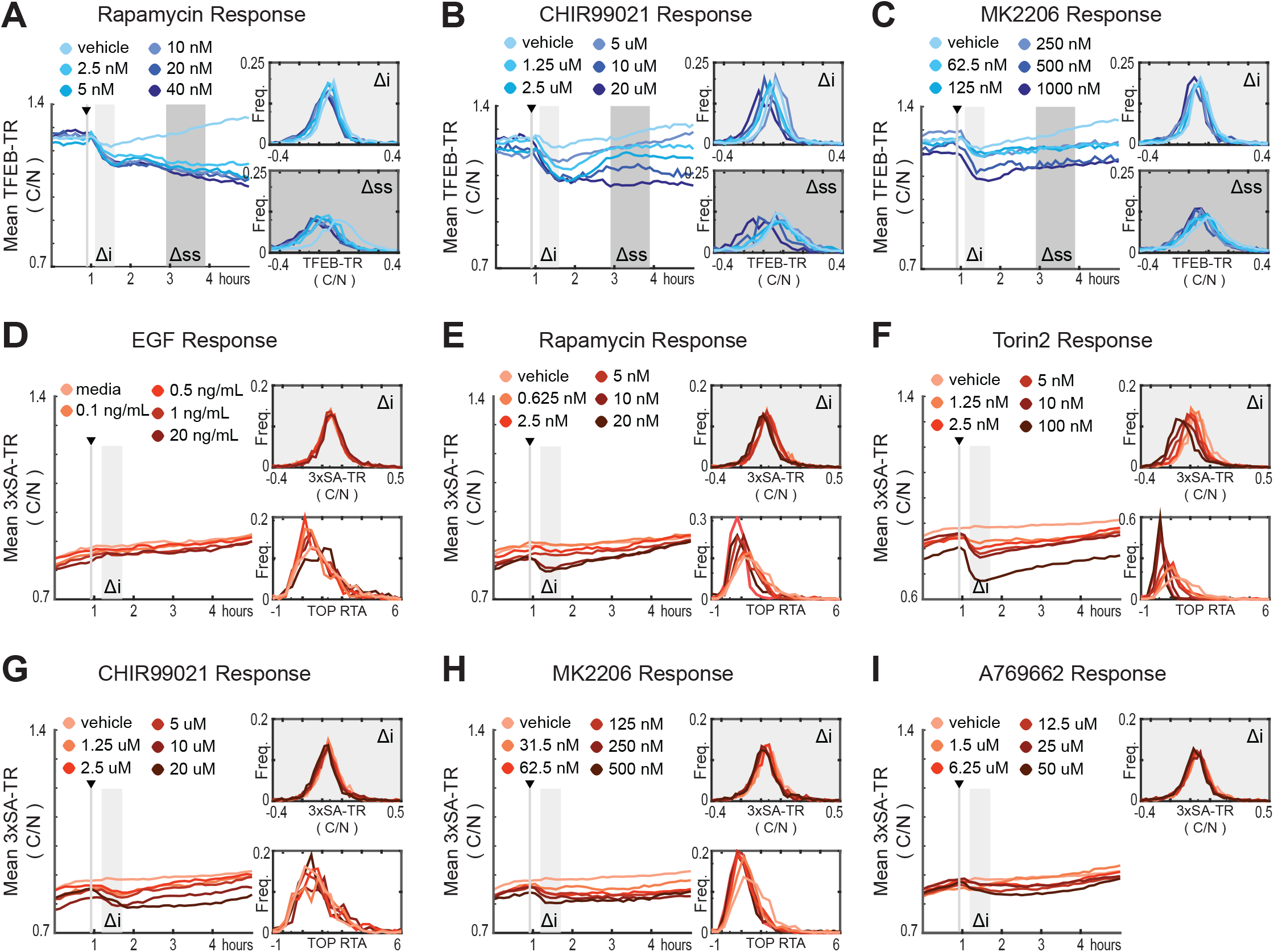
**(A-I)** Quantification of responses of TFEB-TR, 3xSA-TR, or H2B-TOP-DD to titrations of mTORC1 modulators. The mean C/N ratio over time for over >500 cells is reported as a line graph, with histograms of the initial (Δi) and steady state (Δss) responses for individual cells. Cells were starved of growth factors then treated with titrations of (B&H) Rapamycin (A&E), GSKβ inhibitor CHIR99021 (B&G), AKT inhibitor MK2206 (C&H), EGF(D), Torin2 (F), and AMPK activator A769662 (I).

**Figure S5:**
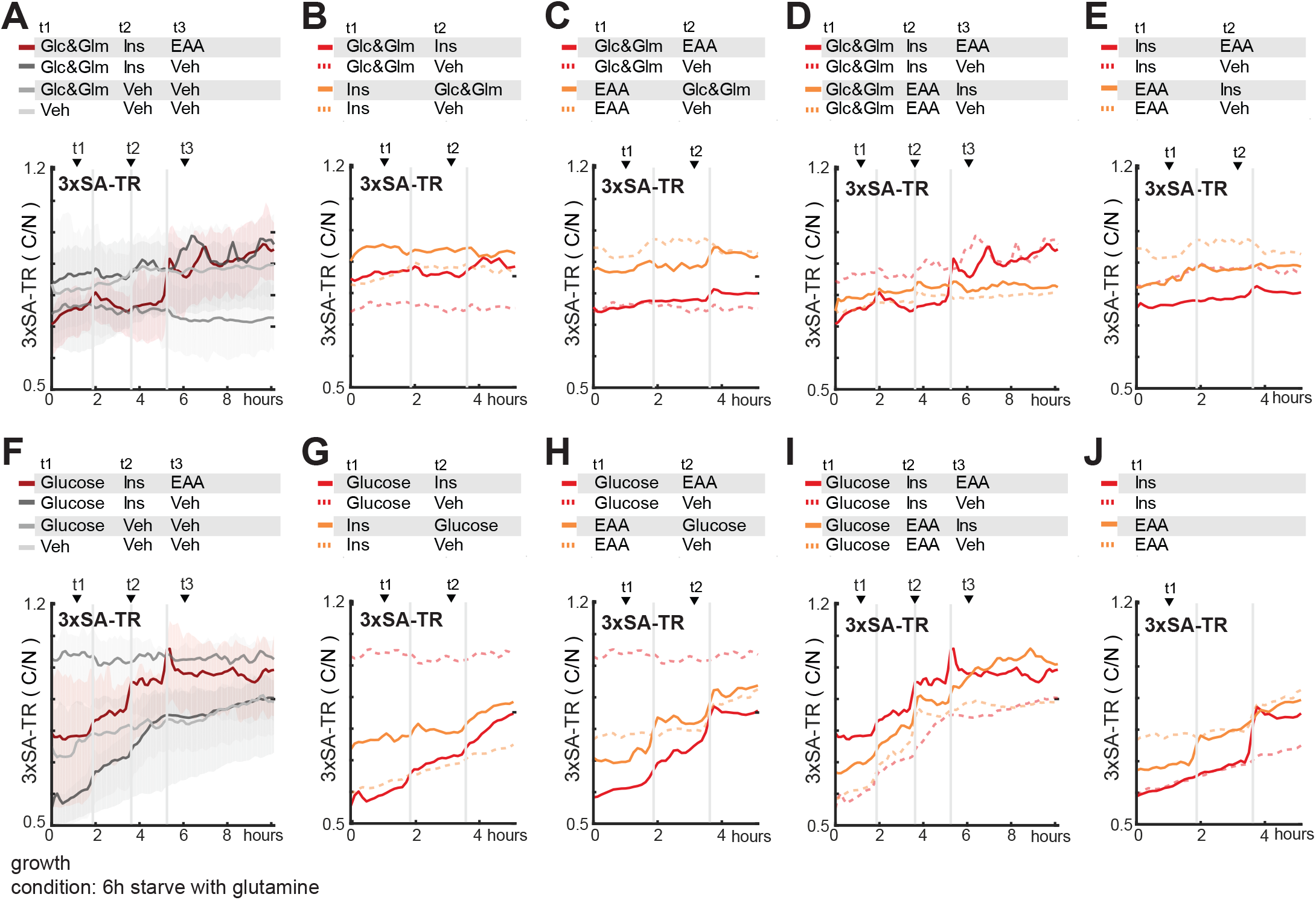
**A-J)** Mean responses of 3xSA-TR to sequential nutrient stimulation in MCF10A cells. In A-E cells were starved of glucose and glutamine, amino acids, and growth factors nutrients for 6 hours, and then stimulated with glucose (17.5mM) and glutamine (2.5 mM), essential amino acids (0.1x), and insulin (10 ug/mL) in the indicated sequences. In F-J cells were starved of glucose, amino acids, and growth factors nutrients for 6 hours, and then stimulated with glucose (17.5mM), essential amino acids (0.1x), and insulin (10 ug/mL) in the indicated sequences. The mean and 25th/75th percentile were calculated from >600 cells. Dotted lines represent media-only controls for each stimulus. Note that the vertical axis limits have been shifted relative to Fig. 3 to better visualize the low dynamic range of 3xSA-TR.

**Figure S6:**
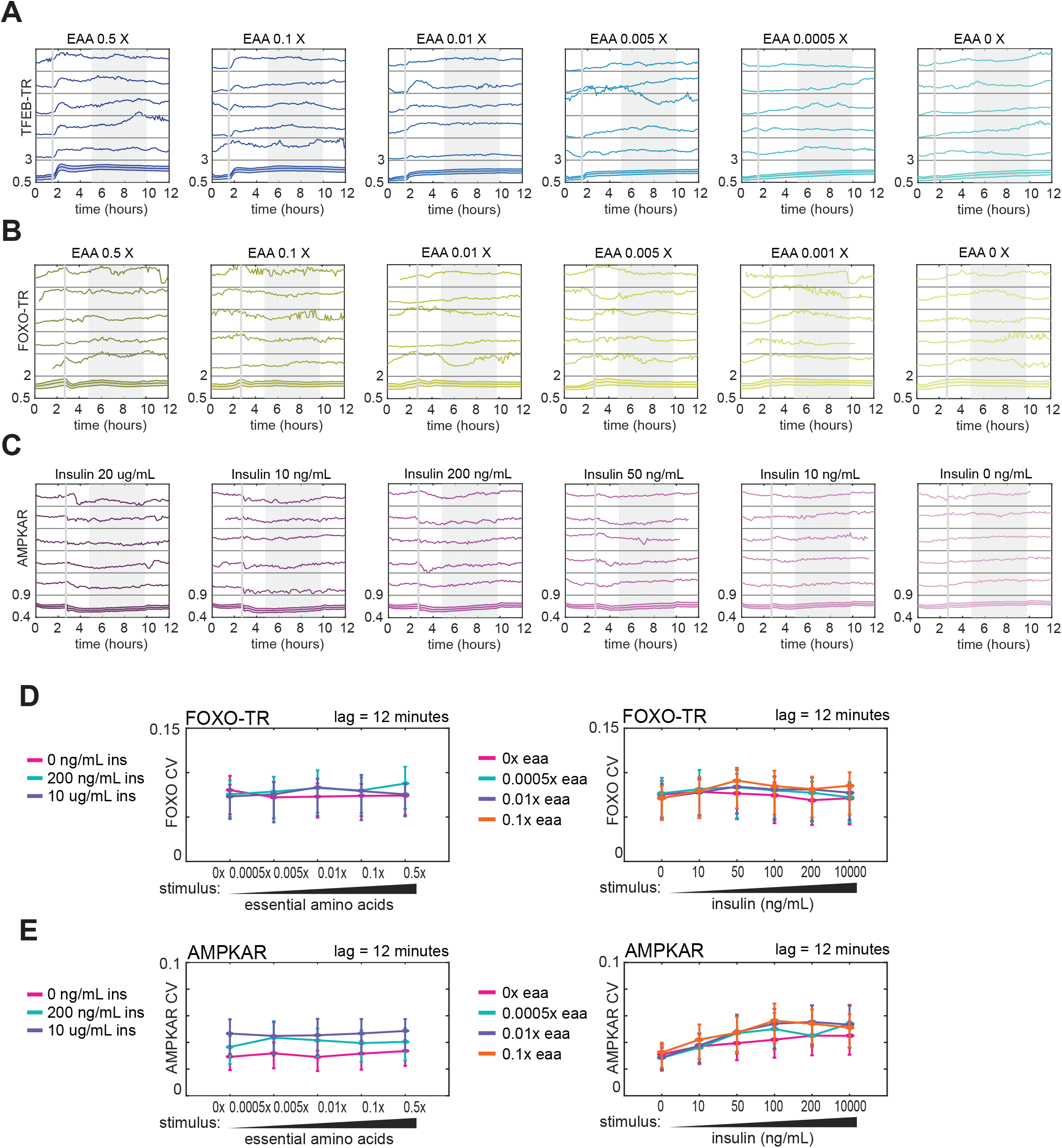
**A)** Single cell TFEB-TR N/C over time. Cells were starved of amino acids for six hours and then stimulated with essential amino acids ranging from 0x to 0.5x concentrated. **B)** Single cell FOXO-TR C/N over time. Cells were grown in a gradient of essential amino acids from 0x to 0.5x (with 2.5mM glutamine, 4mM glucose, and zero growth factors). **C)** Single cell FRET for AMPK activity reporter over time. For six hours, cells were cultured with zero growth factors and 0.1x essential amino acids (with 2.5 mM glutamine), then stimulated with a gradient of insulin. **D)** Single cell, coefficient of variation in FOXO-TR N/C for >400 cells per condition. Reporter expressing MCF10A cells were cultured in gradients of insulin then stimulated with gradients of essential amino acids. Variance calculated as in F6D, with error bars representing the interquartile range. **E)** Single cell, frame-to-frame variance in FOXO-TR N/C for >400 cells per condition. Reporter expressing MCF10A cells were cultured in gradients of amino acids then stimulated with gradients of insulin. Variance calculated as in F6D.

